# Host-Guest binding free energies à la carte: an automated OneOPES protocol

**DOI:** 10.1101/2024.08.23.609378

**Authors:** Pedro Febrer Martinez, Valerio Rizzi, Simone Aureli, Francesco Luigi Gervasio

## Abstract

Estimating absolute binding free energies from molecular simulations is a key step in computer-aided drug design pipelines, but agreement between computational results and experiments is still very inconsistent. Both the accuracy of the computational model and the quality of the statistical sampling contribute to this discrepancy, yet disentangling the two remains a challenge. In this study, we present an automated protocol based on OneOPES, an enhanced sampling method that exploits replica exchange and can accelerate several collective variables, to address the sampling problem. We apply this protocol to 37 host-guest systems. The simplicity of setting up the simulations and of producing well-converged binding free energy estimates without the need to optimize simulation parameters provides a reliable solution to the sampling problem. This, in turn, allows for a systematic force field comparison and ranking according to the correlation between simulations and experiments, which can inform the selection of an appropriate model. The protocol can be readily adapted to test more force field combinations and study more complex protein-ligand systems, where the choice of an appropriate physical model is often based on heuristic considerations rather than a systematic optimization.

## Introduction

The use of computer simulations to estimate the absolute binding free energy between proteins and ligands is a key step in computer-aided drug development^1–4^. The analysis of such simulations provides invaluable insights for understanding protein-ligand binding at the atomistic level and for guiding the design of optimized ligands^5,6^. Optimal ligands typically have a high affinity and a long residence time which, in turn, ensures a sustained pharmacological effect^7,8^. Clearly, a key requirement for the computed binding affinities is that they correlate well with the experimental ones. However, it is often difficult to achieve a reliable agreement between simulations and experiments.^9–11^ Furthermore, it is difficult to systematically improve upon this agreement in the context of the vast and rapidly evolving field of computational techniques and force fields by testing their combinations on large and flexible ligand-protein complexes.

Hence, a number of simplified systems, known as host-guest systems, have been adopted by the community^12^, to provide easier and computationally cheaper benchmarks on which to test computational methodologies that aim to be scaled up to more complex applications. In this context, hosts represent small-size proteins with druggable cavities and guests represent drug-like molecules. When combined, the resulting interactions between the two and the surrounding solvent offer a small scale representation of those occurring in larger scale systems^13–16^. The popularity of such systems surged with the introduction of the SAMPL challenges, starting in 2011 with the SAMPL3 Challenge, in which computational techniques were put to the test to blindly predict a set of physical properties and were later compared to experiments^17–22^.

Limitations that computational techniques typically encounter in correlating to experiments are twofold: the choice of the force field^23^ and the sampling quality^9,24^. The force field has an essential role in determining the simulation’s outcome and includes the choice of a number of factors such as the protein Lennard-Jones parameter, the electrostatic potential for the partial charges, the ligand parameterization and the water model. It is not trivial to establish *a priori* which combinations of parameters would offer the best correlation with experiments for a given system. Likewise, to systematically test a number of models and determine *a posteriori* which one agrees best with experiments is likely system-dependent and often unfeasible because of the high computational cost of exploring a large parameter space on a significant number of systems. Because of these difficulties, in the field it is common practice to pick well-established force fields and models without having methodically determined whether the chosen model is optimal.

The other challenge that limits the accuracy of the free energies derived from the simulations is the quality of the statistical sampling. In principle all the relevant states in a binding process - the most stable binding poses and the unbound solvated state - must be visited multiple times for correctly estimating average quantities such as the binding affinity. However, such extensive sampling is usually not feasible due to the long simulation time required, which can easily reach several milliseconds or more^25^. Many algorithms have been developed to overcome the sampling problem, including alchemical transformation methods^26^ and collective variable (CV) based enhanced sampling methods^27–33^. While the former are the most widely used in the field and benefit from distinct advantages including the relative simplicity of set-up, they also suffer from well known limitations such as a dependency on the chosen binding pose and a difficulty in correctly quantifying key entropic factors such as interfacial water and conformational changes in the target protein.^34–36^.

CV-based methods allow to explicitly sample binding and unbinding events along physical pathways and are being actively developed as an alternative to alchemical approaches that typically employ unphysical paths connecting two end states. CV-based methods are able to explore different binding poses^37,38^ and are less affected by conformational changes of the target and hydration shells, but, to fully converge the free energy profiles, they typically require an optimal choice of CVs finely tuned to each ligand and protein system.^39–43^. For enhanced sampling simulations to achieve an exhaustive sampling in a feasible amount of computer time, the CVs must be able to capture and accelerate all the relevant slow degrees of freedom of the systems. However, the construction of optimal CVs is not trivial and is usually system dependent.^44–49^

In fact, even for small ligands binding to superficial cavities, commonly used CVs based on protein-ligand distances and ligand orientation are often unable to converge the associated free energy profiles in a reasonable simulation time because the sampling of other slow variables such as interfacial water molecules and ligand torsion angles must also be accelerated.^27,37,42,50–54^ For more complex systems, where the binding involves winding paths along curved tunnels and local and non-local conformational changes of the target, the design of optimal CVs is tedious and time consuming to the point that even machine learning approaches need a careful choice of the feature set used in the training to be effective.^32,42^

In this paper, we present an automated protocol that is able to quickly provide reliable binding affinities without the necessity to craft system-dependent CVs. We employ a method that we have recently developed, OneOPES^55^, that takes advantage of different variants of the On-the-fly Probability Enhanced Sampling method (OPES)^56,57^, replica exchange^58^, multi-CV enhanced sampling and a funnel-shaped restraint^59,60^ to extensively sample systems by accelerating relevant slow degrees of freedom. We show that, through the use of multiple replicas and auxiliary CVs, the convergence of the reconstructed free energy profiles is much less dependent on the choice of CVs and the initial binding pose.

We systematically test our protocol on host-guest systems, starting from a subset of 9 host-guest systems from the SAMPL6^20^ and SAMPL8^22^ challenges. The host that we choose - CB8 - is a neutral macrocyclic molecule that is known to be a challenging system that highlights the limits of free energy methodologies^61^. We combine the GAFF2 force field^62^ with three water models - TIP3P^63^, TIP3P-FB^64^ and OPC3^65^ - and three electrostatic models - vacuum, dieletric implicit solvent and AM1-BCC^66^ - and find that one combination, namely GAFF2 with re-fitted dihedral potentials, vacuum charges and TIP3P, clearly offers a better agreement between calculations and experiments, with a good correlation and no evident systematic errors. Using this model, we run independent simulations in triplicate over the entirety of the systems comprising 18 ligands to scrupulously assess the replicability of the results and the quality of the estimates and their errors. We also measure how much our results depend on the availability of a good quality binding pose to start simulations from by comparing binding free energy estimates obtained from optimal starting points with simulations starting from states where the guest is even outside of the binding pocket. Throughout, we provide a number of advanced metrics and their confidence interval to quantitatively validate the results and compare with the best available results.

To assess the protocol’s transferability and to better determine which electrostatic model is preferable in the context of charged systems, we apply it on another set of systems from the SAMPL5^19^, SAMPL6^20^ and SAMPL8^22^ challenges. We choose host TEMOA that is a strongly negatively charged macrocyclic oligomer and 19 guests that are either negatively or positively charged. In this case the dielectric charge model of charges offers a more consistent agreement with experiments. In analogy with the CB8 case, we repeatedly run independent simulations with this model, also starting from different binding poses and evaluate the corresponding advanced metrics.

Overall, we show that applying the OneOPES protocol to a large number of host-guest systems guarantees a high quality of sampling that consistently produces well-converged binding free energy results, disregarding the need for highly optimized CVs or initial binding poses. Our protocol is easily adaptable to other ligand and water force fields and, by carefully comparing computational and experimental results, can guide users in the vexing task of choosing an optimal combination of models. In addition, by providing insight into why one model agrees with experiment better than another, it can also drive the development of improved force fields, so that simulations can be used with increasing confidence in future drug development pipelines.

## Methods

In this section, we analyze in detail the components of the workflow that, starting from the selection of hosts and guests structures, conveniently leads to the estimation of their binding free energies. First, we introduce the OneOPES enhanced sampling technique, then we break down each step of the protocol linking structure selection to OneOPES input preparation. Finally, we discuss computational details and post-processing procedures.

### OneOPES in a Nutshell

OneOPES^55^ is a replica-exchange enhanced sampling scheme that combines different variants of the On-the-fly Probability Enhanced Sampling (OPES)^56,57,67^ method with the aim to effectively accelerate the occurrence of rare events and converge the estimates of free energies even in complex systems, without the need to craft optimized and system-dependent CVs. In OneOPES, one typically sets up 8 simulations in a replica-exchange framework, with replica 0 being the convergence-dedicated one on which equilibrium properties are evaluated and replicas 1-7 being the exploration-dedicated replicas.

All replicas include an OPES Explore^57^ bias that is applied on a set of main CVs. Akin to its predecessor Well-Tempered Metadynamics^68^, OPES Explore makes the system sample a broadened target probability distribution 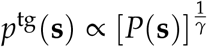 called the Well-Tempered distribution by iteratively building a bias potential *V*(**s**) that, at step *n*, corresponds to

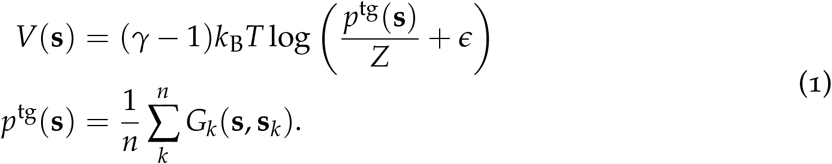

where **s** are the CVs over which the bias is deposited, *γ* = Δ*E*/(*k*_B_*T*) is the bias factor that controls the broadening of the probability distribution, *P*(**s**) is the unbiased marginal distribution, *k*_B_ is the Boltzmann constant, T the temperature given by the thermostat, *Z* a normalization factor, 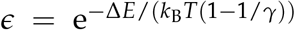 a term that controls the maximum bias deposition and Δ*E* is the corresponding tunable parameter called the `BARRIER`. During the simulation, *p*^tg^(**s**) is estimated on-the-fly by depositing Gaussian kernels such as *G*_*k*_(**s, s**_*k*_) on CVs **s**_*k*_ at the *k*-th step.

In the common situation where the CVs **s** are not able to comprehensively capture all the slow degrees of freedom of the system, the sampling would be hampered and may end up requiring very long simulation times to produce converged free energy results^30,31,39,40^. To alleviate this problem, in OneOPES one introduces a ladder of exchanging replicas that, beside the main OPES Explore bias, include a number of weaker OPES Explore bias potentials over auxiliary CVs and also an OPES Multithermal bias^56^ that, by progressively heating up the system, helps in lowering all kinetic barriers (see Tab. 1).

In the context of ligand binding, standard CVs include the position of the center of mass of the ligand with respect to the binding site and the relative orientation between guest and host. Helpful auxiliary CVs must capture ignored important degrees of freedom such as the hydration of pockets where long-lived water molecules may lie and the guest’s torsional angles.

### An Automated Protocol for System Preparation

The opportunity to efficiently calculate absolute binding free energies with OneOPES allows us to systematically compare various combinations of forcefields and water models. To better handle the data and to facilitate future tests, we propose a protocol that automates system preparation and simulation setup. The corresponding scripts are all available on GitHub at this link and the whole workflow is illustrated in Figure 1.

**Figure 1:**
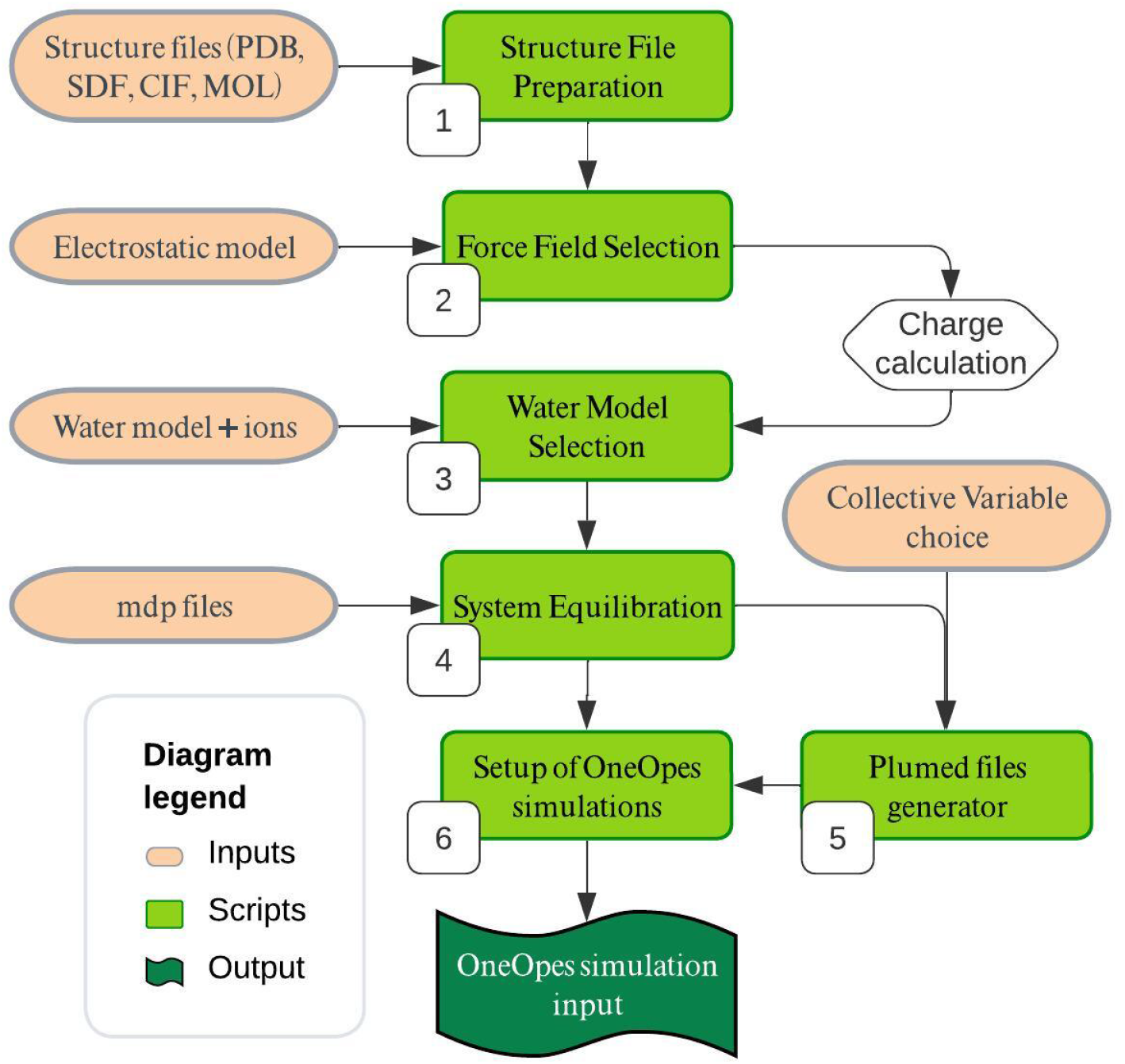
Workflow illustrating each step of the protocol, starting from the structure file preparation to the generation of the OneOPES simulations input.

#### Structure File Preparation and Force Field Selection (1-2)

The protocol begins with the preparation of structure files for the host-guest systems. To favor reproducibility within the Computer-Aided Drug Design (CADD) community, we have implemented support for a variety of standard molecular file formats, including PDB, SDF, CIF, and MOL. This flexibility ensures compatibility with a wide range of input sources and allows for seamless integration into existing workflows.

The following step involves generating a topology of the system under investigation by selecting the appropriate force field for host, guest, and water. While for biological moieties (e.g. proteins or nucleic acids) there are refined force fields already implemented in common MD engines, organic small molecules such as hosts and guests have to be typically parametrized through general force fields such as GAFF2^69^. GAFF2 assigns parameters to each molecule through an atom-typing algorithm that identifies atom types through their chemical environment. Once the atom types are identified, GAFF2 assigns the appropriate force field parameters, including bond lengths, angles, dihedrals, and van der Waals interactions, from its extensive parameter database. Concerns arise when dealing with the electrostatic potential of the overall molecules, which must be converted into atom-centered charges in order to be used in classical MD simulations.

Semi-empirical models are available and largely exploited (e.g. AM1-BCC^66^), ensuring a good balance between computational efficiency and accuracy. Higher-level quantum mechanical methods are able to take into account subtle electronic effects and produce more accurate electrostatic potential-derived charges^70^. To perform these calculations, we employ Gaussian 16^71^. For both the hosts and the guests, we first perform a preliminary energy minimization step using the B3LYP functional^72^, followed by a second minimization step using the Hartree-Fock method^73^ in which single point charges on each atom are calculated via the RESP method^74^. In this step we explore two different options: in one we calculate the electrostatic potential in vacuum, whereas in the other one we use an implicit solvent that mimics the screening effect of an aqueous solution.

Accurately modeling the potential energy curves of rotatable dihedral angles is essential to capture the flexibility of the ligands. Here, we use the AIMNet2 neural network potential,^75^ integrated into Accellera’s Playmolecule^76,77^. Our pipeline can be trivially adapted to use other codes and levels of theory. Through AmberTools’ Parmed2, we convert the coordinates and topology files of each host-guest complex into a format that is compatible with the molecular dynamics engine GROMACS^78,79^.

#### Water Model Selection (3)

The topology files of each individual host and guest are systematically merged into a single file corresponding to each host-guest complex. This unified file must also incorporate additional information such as the selection of the water model and ion parameters. Over the years, numerous water models have been developed to more accurately replicate the properties of bulk water^80–85^. However, one cannot assume that such models can be easily combined with general ligand force fields to capture the behavior of small molecules in aqueous solutions. The choice of water model is particularly critical in this context, not only because water molecules vastly outnumber all other species in a typical biomolecular simulation box, but also because getting the balance right between host-guest interactions and guest and host-water interactions with water is paramount. The choice can significantly impact the calculation of a system’s properties, especially in processes involving hydration or hydrophobic effects.

In the present work, we compare the standard TIP3P water model ^63^, which is widely used in the protein-ligand binding community and was used in the parameterization of GAFF2^62,86^, with two more modern 3-point water models, TIP3P-FB ^64^ and OPC3^65^. Salinity concentrations have been fine-tuned according to the experimental data for each host-guest system. The parameters of the ions used in combination with TIP3P-FB and OPC3 are those reported by Sengupta et al.^87^.

#### System Equilibration (4)

The systems are equilibrated following a standard protocol, 1 ns NVT simulation followed by 2 ns NPT simulation with restraint on heavy atoms. The restraint is then removed and a short 1 ns MD simulation is performed to extract the standard deviations of the CVs to be used in the subsequent OneOPES runs. The equations of motion are integrated with the leap-frog algorithm, using a time step of 2 fs and constraining the hydrogen stretching modes with the LINCS algorithm^88^. The particle-mesh-Ewald (PME) method is used to treat the electrostatic interaction^89^ and a cut-off distance of 1.0 nm is applied on van der Waals interactions. The pressure is set at a reference value of 1 bar with the C-rescale barostat^90^, whereas the temperature is set at 298K with the V-rescale thermostat^91^.

#### Setup of OneOPES Simulations (5-6-7)

The OneOPES simulations are run with the MD engine GROMACS 2023^78,79^, patched with PLUMED2-2.9^92^. To this end, one has to generate PLUMED input files. To limit the guest exploration in the solvated unbound state, we apply a funnel-shaped restraint to confine the guests to a cylindrical volume in the unbound state without affecting the binding path or pose^59^. Given the symmetry of host CB8, we allow the ligand to unbind from two directions and thus include two symmetric funnel restraints, while for host TEMOA we only use one.

On replicas 0-7, we employ as a CV in the main OPES Explore bias the *z* coordinate of the ligand’s center of mass along the funnel’s axis to monitor and accelerate the binding of the guests to their host. Alongside *z*, we also use as a main CV *COS*, the cosine of the angle formed by a ligand axis with the funnel axis. While we do not expect that there are relevant kinetic barriers on *COS*, its use as a main CV helps in distinguishing different poses regarding the orientation of the guest with respect to the host. The parameter SIGMA is derived from earlier unbiased MD simulations, while BARRIER is 100 kJ/mol and PACE 10000 integration steps.

To increase the quality of the sampling, auxiliary CVs are introduced on additional biases OPES MultiCV on replicas 1-7. These include water coordination around selected ligand atoms (*WL*) and on selected points along the funnel axis (*WH*), in analogy with previous work^37,42,93^, and, when present, ligand torsional angles. The water coordination around atom *i* is calculated according to

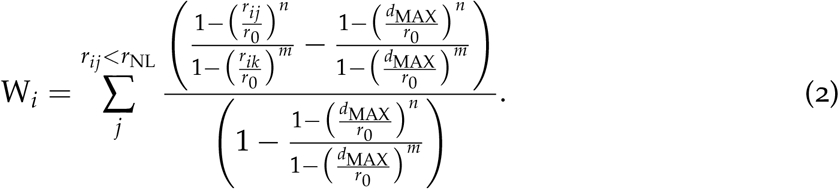

where one sums over all the water oxygen atoms *j, r*_*ij*_ is the distance between central atom *i* and *j, r*_0_ = 2.5 Å is the switching function characteristic distance, *d*_MAX_ = 0.8 Å is a distance at which the switching function smoothly goes to zero, *r*_NL_ = 1.5 Å is the neighbor list cutoff radius and the neighbor list is updated every 20 integration steps. For *WL* we choose *n* = 6 and *m* = 10, while for *WH* we have *n* = 2 and *m* = 6. For these OPES MultiCV biases we use a `BARRIER` of 3 kJ/mol and a PACE of 20000 integration steps.

On replicas 4-7 we also sample increasingly larger temperature ranges via OPES Multithermal with a PACE parameter of 100 integration steps. All these biases are combined in the OneOPES framework, as shown in Tab. 1:

**Table 1:** Schematics of the different CVs and biases used in each replica of the OneOPES simulations. The intersection between rows such as OPES Explore and and columns such as Replica 2 contains the CVs that are biased in that enhanced sampling scheme. If in a replica a bias is not present (e.g. OPES MultiCV 4 and Replica 3), the content of the cell is empty. WL stands for WaterLigand and WH stands for WaterHost. Both indicate the coordination number of the water oxygen atoms with respect to a ligand atom or a virtual atom in the vicinity of the host, respectively. On the row OPES MultiT, we show the highest temperature that we sample in the MultiThermal scheme, with the minimum being the thermostat temperature 298K.

| Replicas | 0 | 1 | 2 | 3 | 4 | 5 | 6 | 7 |
| --- | --- | --- | --- | --- | --- | --- | --- | --- |
| OPES Explore | z, COS | z, COS | z, COS | z, COS | z, COS | z, COS | z, COS | z, COS |
| OPES MultiCV 1 | - | WL<sub>4</sub> | WL<sub>4</sub> | WL<sub>4</sub> | WL<sub>4</sub> | WL<sub>4</sub> | WL<sub>4</sub> | WL<sub>4</sub> |
| OPES MultiCV 2 | - | - | WH<sub>1</sub> | WH<sub>1</sub> | WH<sub>1</sub> | WH<sub>1</sub> | WH<sub>1</sub> | WH<sub>1</sub> |
| OPES MultiCV 3 | - | - | - | WL<sub>1</sub> | WL<sub>1</sub> | WL<sub>1</sub> | WL<sub>1</sub> | WL<sub>1</sub> |
| OPES MultiCV 4 | - | - | - | - | WH<sub>3</sub> | WH<sub>3</sub> | WH<sub>3</sub> | WH<sub>3</sub> |
| OPES MultiCV 5 | - | - | - | - | - | WH<sub>8</sub> | WH<sub>8</sub> | WH<sub>8</sub> |
| OPES MultiCV 6 | - | - | - | - | - | - | WH<sub>5</sub> | WH<sub>5</sub> |
| OPES MultiCV 7 | - | - | - | - | - | - | - | WH<sub>10</sub> |
| OPES MultiT | - | - | - | - | 310K | 330K | 350K | 370K |

When torsional angles associated with slow transitions are present, they are included in all the OPES MultiCV biases. If up to two torsional angles are selected, the OPES MultiCV would include up to 3 CVs - an hydration CV as in Tab. 1 and the torsional angles. In the case of 3 relevant torsional angles, we cycle through them so that each OPES MultiCV bias includes 3 CVs, with different combination of couple of torsional angles present. The number of torsional angles that we selected for each system is reported in Tab. S1 in the Supporting Information (SI).

Exchanges between replicas are attempted every 1000 integration steps. Regarding the thermostat and the barostat, we use the same ones described in the System equilibration. To ensure a swift, error-free setup process, our protocol takes care of simulation input preparation by generating all the required GROMACS and PLUMED input files in the dedicated directories pertaining to each replica.

### Computational Details and Post-processing

In this work we investigate two host systems, i.e. Cucurbit[8]uril (CB8) and tetraendo-methyl-octa-acid (TEMOA), in combination with a number of guest molecules from the SAMPL5^19^, SAMPL6^20^ and SAMPL8^22^ challenges. CB8 is characterized by a symmetrical, circular, and neutrally charged structure, whereas TEMOA is a strongly negatively charged calixarene with a total charge of −8. For a full list of the guest/host combinations that we study, please refer to Figs. 2-3 and Tab. S1. For clarity, we follow a naming convention by calling each guest SX-GY for each host, where X identifies the SAMPL challenge from which the system is taken and Y indicates the corresponding name of the guest. The initial state of the simulations is chosen from the crystal structure of the bound state, when available. To verify that our results do not depend on the chosen initial state, we also test starting conditions that are far from the binding pocket (see Figs. S5-S9).

**Figure 2:**
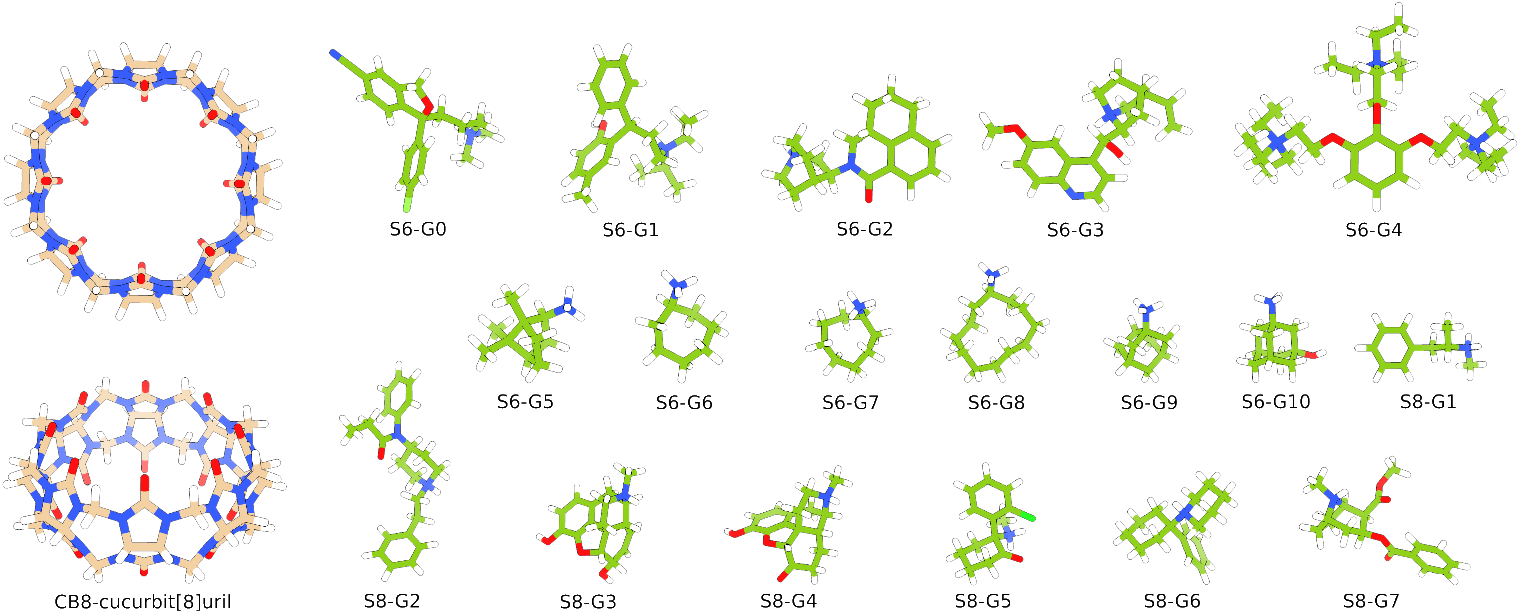
CB8 and guest structures from the SAMPL6 and the SAMPL8 challenges. The subset of ligands S6-G3, S6-G5, S6-G7, S6-G8, S6-G9, S8-G1, S8-G2, S8-G4 and S8-G6 is used in the force field search phase. SAMPL8 ligands correspond to drugs of abuse.

**Figure 3:**
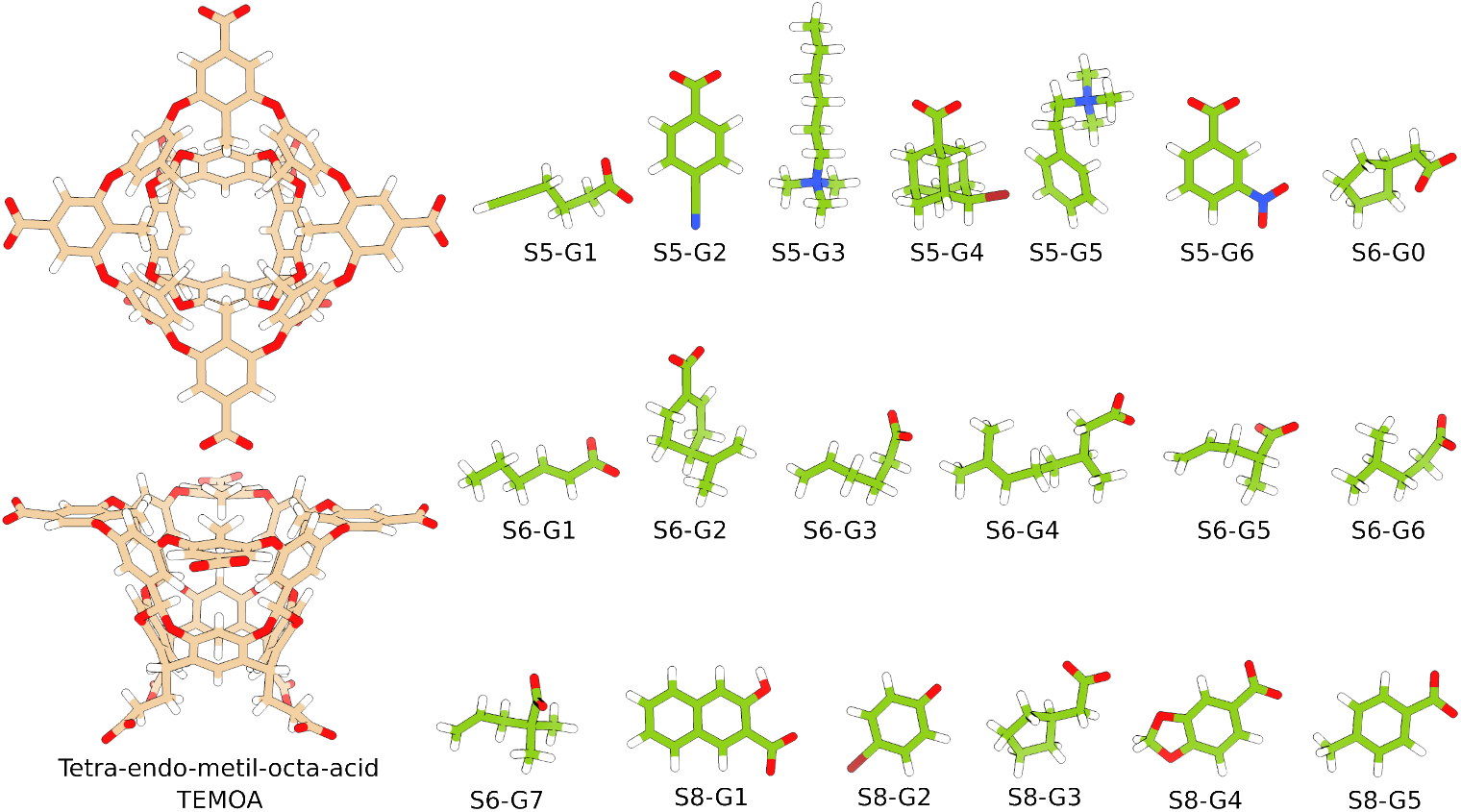
TEMOA and guest structures from the SAMPL5, the SAMPL6 and the SAMPL8 challenges.

Our simulation approach is divided into two main phases:

- **Force Field Search**: we test a selection of guests using different electrostatic options (AM1-BCC, Vacuum, Dielectric) and water models (TIP3P, TIP3P-FB, OPC3) to identify their optimal combination with respect to error and correlation with experimental results. In this case, to obtain a quick binding free energy estimate is preferred and simulations are run in single copy, with the error being evaluated by a block average. This phase helps in establishing a reliable computational framework for the subsequent set of more in-depth simulations.
- **Results Refinement**: we take the optimal force-field from the previous step and study all the available host-guest combinations available - 18 systems for CB8 and 19 for TEMOA. Each OneOPES simulation is run in triplicate to ensure robustness and reproducibility of the results and we also simulate different initial binding conditions.

The use of a funnel-shaped restraint to confine the guests to a cylindrical volume in the solvent introduces an undersampling of the unbound state that must be taken into account when calculating absolute binding free energies. To factor out its effect^28^, the absolute binding free energy must be calculated by applying

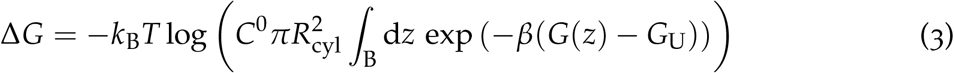

on replica 0, where *k*_B_ is Boltzmann’s constant, *T* the thermostat’s temperature, *C*^0^ = 1/1660 Å^−3^ the standard concentration, *R*_cyl_ = 2 Å the funnel’s cylinder radius, *z* the coordinate of the ligand’s center of mass along the funnel’s axis, *G*(*z*) is the free energy value along the funnel axis and *G*_U_ its reference value in the unbound state.

The symmetrical nature of CB8 and its openness from both ends, encouraged us to include two symmetric funnel restraints, letting the guests unbind from both directions. This choice helps in capturing binding and unbinding events in both directions and in better describing the rich binding pose landscape that can arise from the host’s symmetry. As visible for example in Fig. S14, the apparent symmetry of the resulting free energy profiles is not imposed and corresponds perfectly to the symmetry of the host molecule. This, in turn, helps in highlighting how well simulations converge. The two binding free energy estimates that are generated through Eq. 3 on the two unbound states - above and below the host - are both used in the binding free energy estimation and its error. In the case of large ligands such as S6-G4, when passing through the host, the guest is sterically impeded (see Fig. S5) and one of the unbound states is barely ever visited (see Fig. S17), we only use the binding free energy estimate from the well sampled side.

Errors on the binding free energy estimates are calculated in two different ways. In the force field search phase, after discarding the simulation’s begininning - 50 ns for TEMOA and 100 ns for CB8 - that is considered as equilibration, we divide the remaining portion of simulation in 3 blocks and calculate the system’s free energy over *z* through standard reweighting and its error as the standard deviation over the blocks. In CB8, the presence of the two funnel restraints encourages us to first take the average between the two free energy estimates for each block and then to perform the block average. In the Results Refinement phase, we perform an analogous strategy with the difference that each of the 3 independent simulations that we run represents a block in the block average procedure. To evaluate the quality of the results with respect to experiments, we calculate standard metrics such as Kendall’s *τ*, the linear fit *R*^2^ and slope *m*, the Mean Absolute Error (MAE), the Mean Error (ME), and the Root Mean Squared Error (RMSE). The confidence intervals that we report are calculated via a bootstrap procedure adapted from the one published by Rizzi et al.^61^ where we perform 100000 iterations. All post-processing scripts are available in the GitHub repository https://github.com/Pefema/OneOpes_protocol.

## Results

We first apply the strategy to the neutral host CB8 and then to the negatively charged TEMOA. In both cases, a force field exploration phase, in which we try out a number of force field combinations, is followed by a results refinement phase, in which we select a force field and perform three independent calculations, also starting from different binding poses.

### CB8 Results

#### Force Field Search for CB8

We explore all the combinations of three electrostatic models (AM1-BCC, vacuum and dielectric) with three water models (TIP3P, TIP3P-FB and OPC3) on a a subset of 9 guests and host CB8. This comprehensive approach allows us to evaluate which force field combination better agrees with experimental data. Our results, illustrated in Fig. 4, show a rather clear performance hierarchy among the tested models. The TIP3P water model consistently outperforms the TIP3P-FB and OPC3 across all charge types. Regarding the electrostatics, the vacuum one gives the best agreement with experiment, closely followed by the dielectric model, with the AM1-BCC model showing the poorest performance. In all cases, our results correlate well with experiments, but display different systematic errors. The ability to systematically pinpoint the shortcomings of force fields is one of the original aims of the SAMPL challenges and these results highlights the quality of the sampling granted by our strategy.

**Figure 4:**
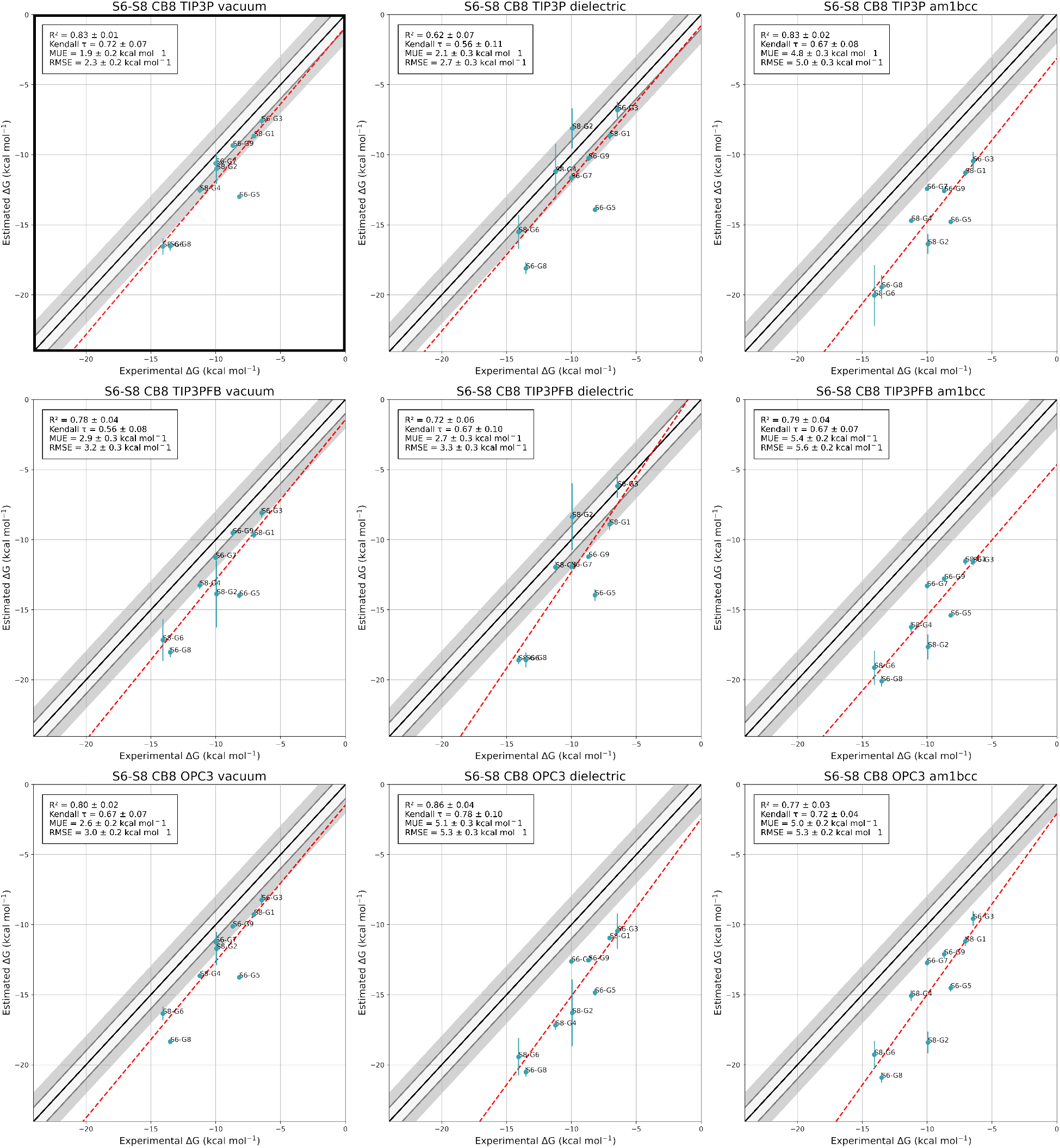
Experimental vs Predicted binding free energies for CB8 for 9 starting ligands with different force field combinations. The rows correspond to TIP3P, TIP3PFB and OPC3 water models and the columns correspond to vacuum, dielectric and AM1-BCC electrostatic models, respectively. The legend shows *R*^2^, Kendall *τ*, Mean Unsigned Error (MUE) and Root Mean Squared Error (RMSE). The combination that shows the best agreement with experiments according to *R*^2^ and RMSE is TIP3P water with vacuum electrostatics and is highlighted in the top left corner.

The combination of TIP3P water with vacuum electrostatic model emerges as the optimal choice, giving the highest correlation with experimental data (*R*^2^ = 0.83) and the lowest root mean square error (RMSE = 2.3 kcal/mol) with respect to experiment. Hence, this optimal model is carried over to the following more extensive set of simulations.

#### Results Refinement for CB8

We extend our analysis to the entire set of 18 CB8 ligands from the SAMPL6 and SAMPL8 challenges. Each simulation is performed in triplicate to highlight the robustness of the estimates and retrospectively validate the cheaper force field search estimations. The results presented in Fig. 5 show a good correlation between experimental and predicted binding free energies (*R*^2^ = 0.69, *m* = 1.11 and Kendall *τ* = 0.62). The RMSE is 2.05 kcal/mol, the MAE 1.73 kcal/mol and the ME of 1.17 kcal/mol. The estimates are consistent across the triplicate runs and all the systems, with an average standard deviation of 0.49 kcal/mol. In Figs. S1-S2 we juxtapose the advanced metrics of our results with those from some of the best submissions from the corresponding SAMPL challenges. Our results consistently rank among the best for each metric that we estimate.

**Figure 5:**
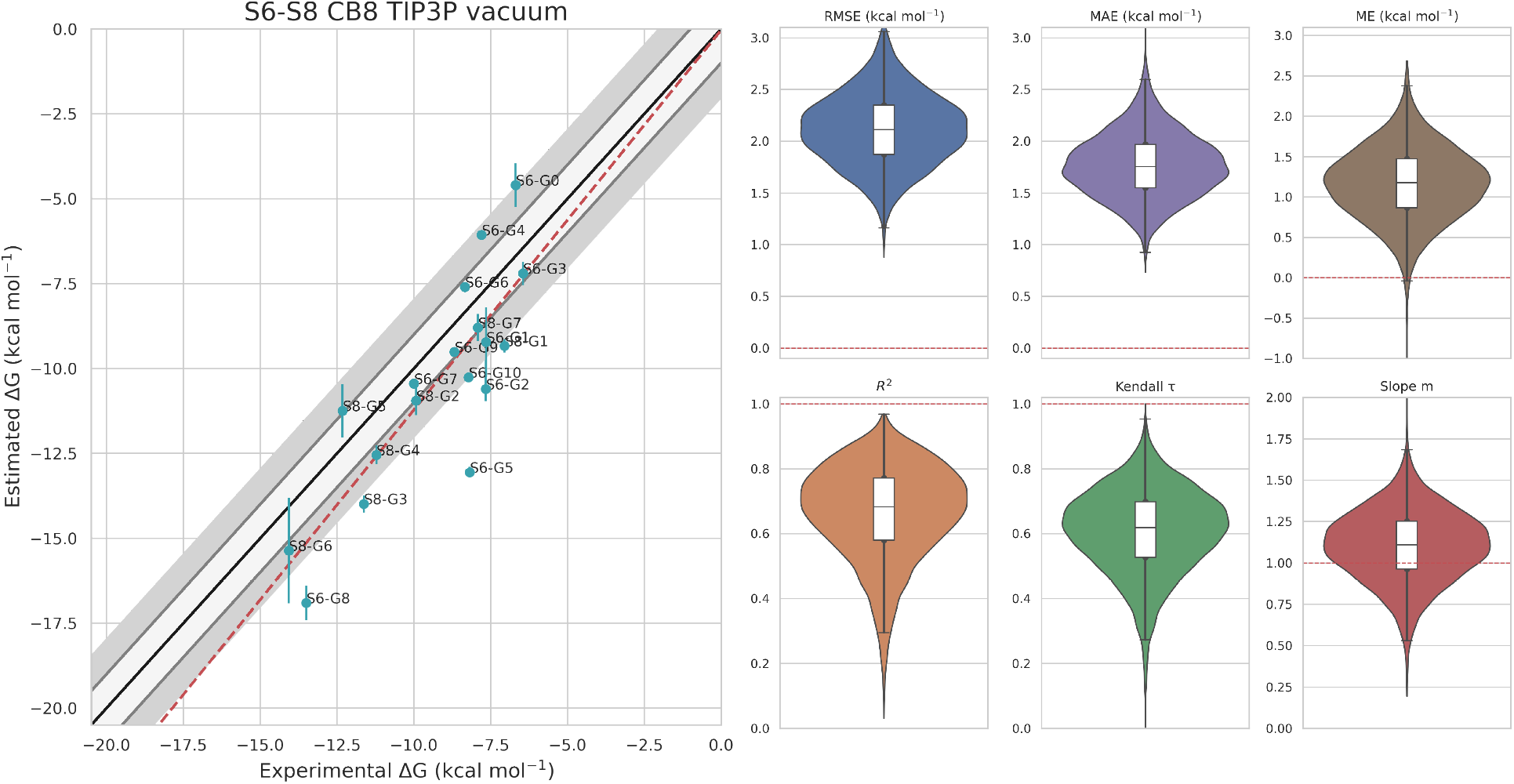
Experimental vs Predicted binding free energies for all 18 CB8 ligands in the Results Refinement phase. All simulations are run in triplicate and we show their average binding free energy and its standard deviation. We show advanced metrics (RMSE, MAE, ME, *R*^2^, Kendall *τ* and *m*) and their confidence interval calculated with a bootstrap technique. The red dashed lines in the boxplots represents the ideal value for each metric.

When comparing the results from the triplicate simulations with those from the single-copy runs from the previous phase (see Fig. S10), we observe a strong correlation between the two with a *R*^2^ = 0.92. This underscores that within our strategy also single-copy simulations provide reliable estimates, closely aligning with the more robust triplicate runs. These results suggest that, while triplicate runs offer increased confidence in the results, especially for more complex ligands, single-copy simulation can often provide a decent enough quality estimate of binding free energies, striking a balance between computational efficiency and reliability. This feature would be particularly endearing in the context of high-throughput screening.

It is not surprising that larger and more flexible ligands such as S6-G1 and S8-G2 exhibit a higher uncertainty in their binding free energy estimations. In particular, the large size of S6-G4 severely hinders the sampling of states that require the guest to squeeze through the host. Because of this difficulty, when one side is severely undersampled, we consider only the sampled side in the binding free energy estimate. Adding torsional angles to the additional CVs that are biased in the strategy offers some help in passing through, but it is clear that, for complex cases such as these to reach a strong convergence in the given simulation time, the ideal solution is to develop ad-hoc CVs that encode all the relevant degrees of freedom. While this route is achievable in a number of ways^45,48,49^, it goes beyond the scope of this paper where our aim is to present a general strategy to build an ensemble view on a whole set of ligands rather than focus on a particular case.

Another crucial aspect for a binding free energy calculation method is its independence from the initial binding pose. Examples are known where competing binding poses both contribute to the experimentally measured binding^94^. In the particular case of CB8, this aspect is even more evident because of the host’s symmetry that lets ligands bind from two opposite directions and makes the presence of multiple binding poses very frequent (see Fig. S14, for example). In the recent work by Karrenbrock et al.^36^, OneOpes was shown not to significantly depend on the starting conformation in protein-ligand systems such as HSP90. Here, we compare triplicate runs starting from the original binding pose provided by the SAMPL challenge with another set of triplicate runs starting from a set of drastically different poses (see Figs. S5-S6), deliberately chosen to be out of the host’s binding site.

As shown in Fig. S11, the two sets of simulations strongly correlate well with each other with a *R*^2^ = 0.91 between the two binding free energies estimates. Remarkably, both sets of simulations converge to comparable results, with only a few cases with larger error bars that correspond to the larger and more flexible ligands. This observation suggests again that more complex system either require longer simulation times or more refined CVs for achieving comparable quality estimates with the smaller ligands. Nevertheless, the overall strong correlation proves the method’s ability to discover more stable binding poses and to identify the correct absolute binding free energy for most compounds, even when starting in poses that are not free energy minima.

### TEMOA Results

#### Force Field Search for TEMOA

To test the method’s transferability, we apply it on TEMOA, a host with a different shape and a different total charge than CB8. TEMOA has appeared in the SAMPL5, SAMPL6 and SAMPL8 challenges is coupled with a total of 19 guests. Given the better performance of the TIP3P water model in the CB8 simulations, we restrict the force field Search phase to that water model. Due to the faster convergence observed in these systems, all simulations are limited to 150 ns per replica.

In this case, the best agreement with experiment is achieved with the dielectric electrostatic model that yields a *R*^2^ = 0.69, *τ* = 0.66, RMSE of 1.3 kcal/mol and MAE of 1.0 kcal/mol (see Fig. 6). Curiously, we observe a striking difference in the response of positively and negatively charged ligands to changes in the electrostatic model. In going from the vacuum to the dielectric model, the estimated binding free energy of all ligands systematically decreases, but the rate of decrease strongly depends on the guests’ charge. For negatively charged ligands we observe a decrease by an average value of 1.5 kcal/mol, while positively charged ligands decrease on average by 3.2 kcal/mol. This observation suggests a systematic charge-dependent role of the electrostatic model in host-guest binding, warranting further investigations.

**Figure 6:**
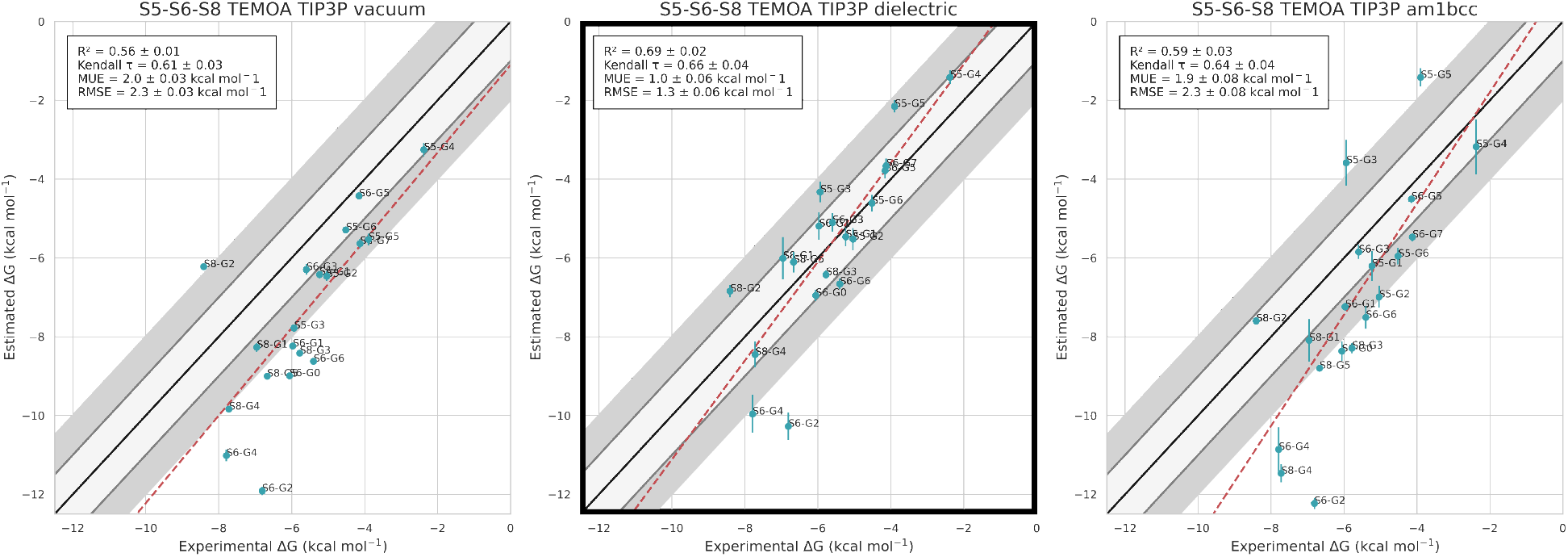
Experimental vs Predicted binding free energies for all 19 TEMOA ligands with different force field combinations. All simulations use the TIP3P water model and the columns correspond to vacuum, dielectric and AM1-BCC electrostatic models, respectively. The legend shows *R*^2^, Kendall *τ*, Mean Unsigned Error (MUE) and Root Mean Squared Error (RMSE). The combination that shows the best agreement with experiments according to *R*^2^ and RMSE is TIP3P water with dielectric electrostatics and is highlighted.

**Figure 7:**
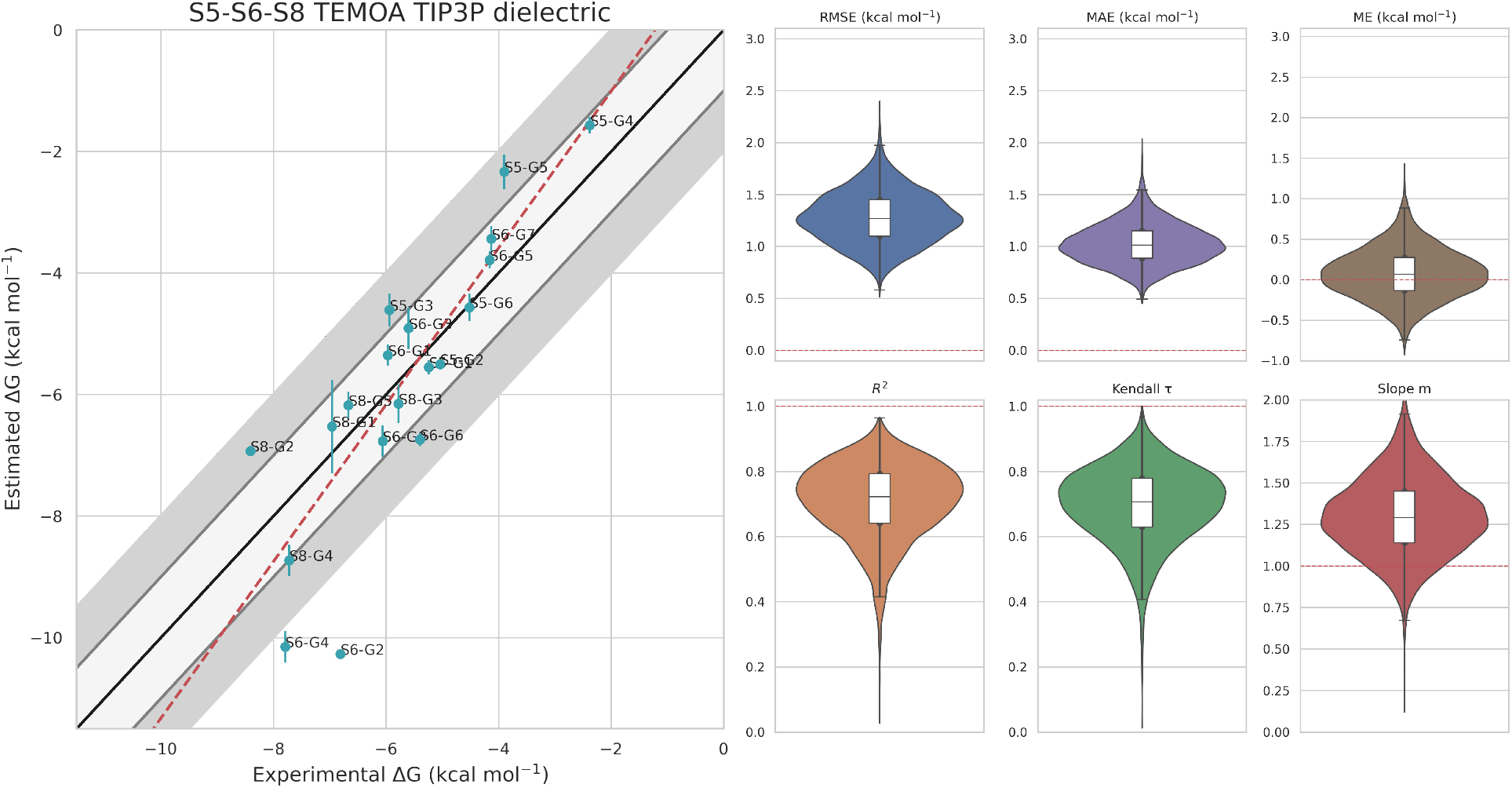
Experimental vs Predicted binding free energies for all 19 TEMOA ligands in the Results Refinement phase. All simulations are run in triplicate and we show their average binding free energy and its standard deviation. We show advanced metrics (RMSE, MAE, ME, *R*^2^, Kendall *τ* and *m*) and their confidence interval calculated with a bootstrap technique. The red dashed lines in the boxplots represents the ideal value for each metric.

#### Results Refinement for TEMOA

In analogy with the CB8 case, we pick the model that best agrees with experiments - TIP3P water and dielectric electrostatics - and perform simulations in triplicate copy. The results, illustrated in Figure 5, show a strong correlation with experiments, with RMSE of 1.26 kcal/mol, MAE of 0.98 kcal/mol, ME of 0.08 kca/mol, *R*^2^ = 0.72, *τ* = 0.73 and *m* = 1.29. A comparison with a selection of the best SAMPL challenge results is shown in Figs. S3-S4. This phase’s results correlate well again with the single-run from the force field search phase, having a *R*^2^ = 0.99 (see Fig. S12). Moreover, also in this case, we repeat simulations starting from significantly different initial states (see Figs. S7-S9) and observe well correlated results with the simulations starting from the given binding poses with a *R*^2^ = 0.94 (see Fig. S13).

## Discussion

We sought to address the challenge of achieving computational binding affinities that closely align with experimental values for host-guest systems by using our recently developed OneOPES free energy approach. We tested different electrostatic potential options and water models, ultimately determining that, with the GAFF2 ligand force field, the standard TIP3P water model generally outperforms more refined 3-point water models such as TIP3P-FB and OPC3. While this result may seem counterintuitive at first, it becomes more clear when considering that the GAFF2 force field was specifically optimized to work with the TIP3P water model^62^. One might even speculate that the superior performance of the GAFF2/TIP3P combination could be due to a cancellation of errors.

With respect to the optimal choice of the point charges, we examined two systems that are characterized by vastly different electrostatic profiles, with CB8 being neutral and TEMOA carrying a significant negative charge. Our findings suggest that for systems like CB8, where the overall electrostatic charge is null, employing vacuum-fitted charges is the optimal approach. Conversely, for systems like TEMOA that exhibit a pronounced total electrostatic charge, charges fitted in the presence of an implicit water model proved to be more effective. This distinction highlights the importance of tailoring electrostatic potential choices to the specific characteristics of the system under study, ensuring the accuracy of the binding affinity predictions. We are planning to further explore the optimal force field search by testing the effect of other options such as polarizable force fields.

Our study’s insights were enabled by the OneOPES enhanced sampling scheme, which, allows an efficient exploration of the host-guest conformational landscape and reliable convergence of the free energy profiles. Remarkably, the excellent agreement between the calculated and experimental binding free energies is achieved independently of the chosen initial binding position. In fact, excellent results were obtained even when the guest molecules were initially placed far away from the host system. The recent application of an equivalent OneOPES strategy on a number of complex protein-ligand systems remarkably shows analogous results^38^. Promising future directions include applying the protocol to systems where the host flexibility plays a role in the binding process or where there are multiple conformations of the binding pocket. The strategy can be trivially adapted to include such degrees of freedom in the CVs space to be enhanced.

It should be emphasized that embedding OneOPES in an automated workflow greatly simplifies the setup of free energy calculations, providing well converged and reproducible free energies that can be used to evaluate novel water models and ligand force fields. In conclusion, our automated OneOPES protocol marks a significant advance in CV-based computational binding free energy estimation. By eliminating the need for system-specific CV definitions and the knowledge of the crystallographic binding poses to converge the binding free energies, it provides a systematic tool for producing accurate and reproducible free energy calculations with minimal user input. Our intention is to further refine this protocol based on feedback from the community and make it a useful tool for improving the accuracy and reproducibility of computational drug discovery pipelines.

## Acknowledgement

The authors acknowledge the Swiss National Science Foundation and Bridge for financial support (projects number: 200021_204795, CRSII5_216587 and 40B2-0_203628) and PRACE and the Swiss National Supercomputing Centre (CSCS) for large supercomputer time allocations on Piz Daint, project IDs: s1169, s1228 and s1274. The authors are grateful to Ioannis Galdadas, Maurice Karrenbrock and Alberto Borsatto for fruitful discussions and to Nicola Piasentin for carefully reading the manuscript.

## Supporting Information Available

Supporting information: Additional figures, tables and computational details. The GRO-MACS and PLUMED input files and the protocol scripts are all available on GitHub at: https://github.com/Pefema/OneOpes_protocol and on PLUMED NEST^95^.

## TOC Graphic

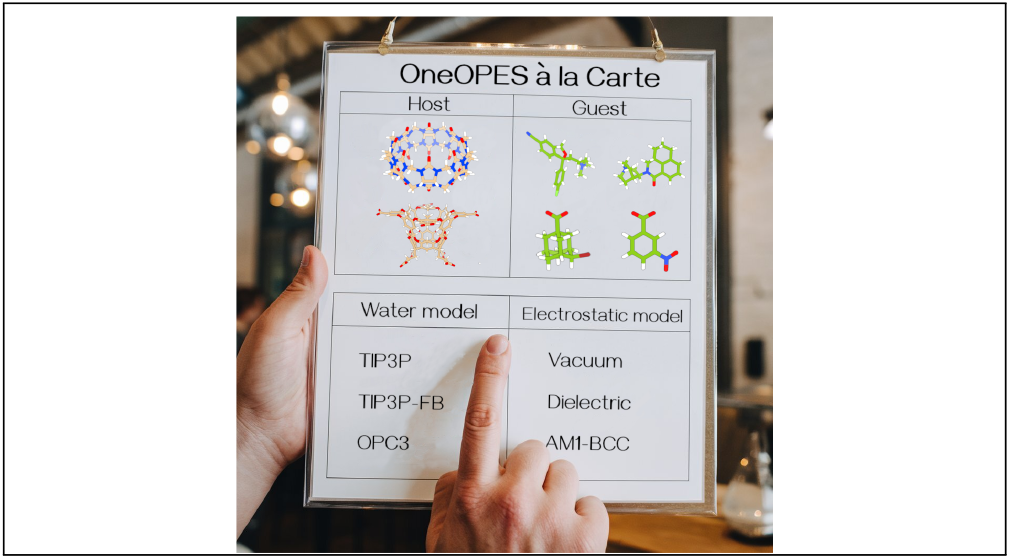

## Notes

### Competing Interest Statement

The authors have declared no competing interest.

### Summary of Updates

There are improvements in the text, new simulations for TEMOA with the AM1-BCC electrostatic model and updated figures.

