## Supporting Information for "Host-Guest binding free energies à la carte: an automated OneOPES protocol"

**Table S1:** Table containing relevant information about all the host-guest systems that we simulate, including the ligands' name, their experimental and OneOPES simulated binding affinities, our simulation time and the number of torsional angles biased.

| Host-Guest | $\Delta G_{\mathrm{OneOPES}}$ (kcal/mol) | $\Delta G_{\mathrm{exp}}$ (kcal/mol) | ns/replica | N° Biased torsions | Compound Name |
|---|---|---|---|---|---|
| CB8-S6-G0 [1] | -4.60 ± 0.64 | -6.69 | 200 | 3 | escitalopram |
| CB8-S6-G1 [1] | -9.53 ± 1.50 | -7.65 | 200 | 3 | tolterodine |
| CB8-S6-G2 [1] | -10.61 ± 0.36 | -7.66 | 200 | - | palonosetron |
| CB8-S6-G3 [1] | -7.43 ± 0.65 | -6.45 | 200 | 3 | quinine |
| CB8-S6-G4 [1] | -6.07 ± 0.08 | -7.80 | 200 | 3 | gallamine triethiodate |
| CB8-S6-G5 [1] | -13.06 ± 0.16 | -8.18 | 200 | - | (1R,2S,4R)-1,7,7-trimethylbicyclo[2.2.1]heptan-2-amine |
| CB8-S6-G6 [1] | -7.60 ± 0.14 | -8.34 | 200 | - | cycloheptanamine |
| CB8-S6-G7 [1] | -10.45 ± 0.12 | -10.00 | 200 | - | cyclododecanamine |
| CB8-S6-G8 [1] | -16.91 ± 0.51 | -13.50 | 200 | - | cyclooctanaine |
| CB8-S6-G9 [1] | -9.51 ± 0.13 | -8.68 | 200 | - | (2R,3as,5S,6as)-hexahydro-2,5-methanopentalen-3a(1H)-amine |
| CB8-S6-G10 [1] | -10.27 ± 0.14 | -8.22 | 200 | - | (1S,3r,5R,7S)-3-aminoadamantan-1-ol |
| CB8-S8-G1 [2] | -9.34 ± 0.19 | -7.05 | 200 | 1 | Meth |
| CB8-S8-G2 [2] | -11.16 ± 0.97 | -9.93 | 200 | 3 | Fentanyl |
| CB8-S8-G3 [2] | -13.99 ± 0.26 | -11.63 | 200 | - | Morphine |
| CB8-S8-G4 [2] | -12.55 ± 0.28 | -11.22 | 200 | - | Hydromorphone |
| CB8-S8-G5 [2] | -11.24 ± 0.79 | -12.32 | 200 | 2 | Ketamine |
| CB8-S8-G6 [2] | -15.36 ± 1.55 | -14.07 | 200 | 2 | PCP |
| CB8-S8-G7 [2] | -8.80 ± 0.40 | -7.92 | 200 | 3 | Cocaine |
| TEMOA-S5-G1 [3] | -5.55 ± 0.03 | -5.24 | 150 | - | 5-Hexenoic acid |
| TEMOA-S5-G2 [3] | -5.50 ± 0.00 | -5.04 | 150 | - | 4-Cyanobenzoic acid |
| TEMOA-S5-G3 [3] | -4.61 ± 0.06 | -5.94 | 150 | - | Hexyltrimethylammonium |
| TEMOA-S5-G4 [3] | -1.57 ± 0.03 | -2.38 | 150 | - | 4-bromoadamantane-1-carboxylic acid |
| TEMOA-S5-G5 [3] | -2.33 ± 0.07 | -3.90 | 150 | - | trimethyl(2-phenylethyl)azanium |
| TEMOA-S5-G6 [3] | -4.56 ± 0.05 | -4.52 | 150 | - | 3-nitrobenzoic acid |
| TEMOA-S6-G0 [1] | -6.77 ± 0.06 | -6.06 | 150 | - | cyclopentyl acetic acid |
| TEMOA-S6-G1 [1] | -5.35 ± 0.04 | -5.97 | 150 | - | trans-2-hexenoic acid |
| TEMOA-S6-G2 [1] | -10.27 ± 0.00 | -6.81 | 150 | - | (s)-(-)-perillic acid |
| TEMOA-S6-G3 [1] | -4.91 ± 0.08 | -5.60 | 150 | - | 5-hexanoic acid |
| TEMOA-S6-G4 [1] | -10.15 ± 0.06 | -7.79 | 150 | - | (s)-(-)-citronellic acid |
| TEMOA-S6-G5 [1] | -3.79 ± 0.03 | -4.16 | 150 | - | 2-methyl-4-pentenoic acid |
| TEMOA-S6-G6 [1] | -6.74 ± 0.03 | -5.40 | 150 | - | 4-methylpentanoic acid |
| TEMOA-S6-G7 [1] | -3.44 ± 0.05 | -4.13 | 150 | - | 2,2-dimethyl-4-pentenoic acid |
| TEMOA-S8-G1 [2] | -6.53 ± 0.18 | -6.96 | 150 | - | 3-hydroxy-2-naphthoic acid |
| TEMOA-S8-G2 [2] | -6.93 ± 0.02 | -8.41 | 150 | - | p-bromophenol |
| TEMOA-S8-G3 [2] | -6.15 ± 0.08 | -5.78 | 150 | - | cyclopentylacetic acid |
| TEMOA-S8-G4 [2] | -8.72 ± 0.06 | -7.72 | 150 | - | piperonylic acid |
| TEMOA-S8-G5 [2] | -6.17 ± 0.05 | -6.67 | 150 | - | p-toluic acid |

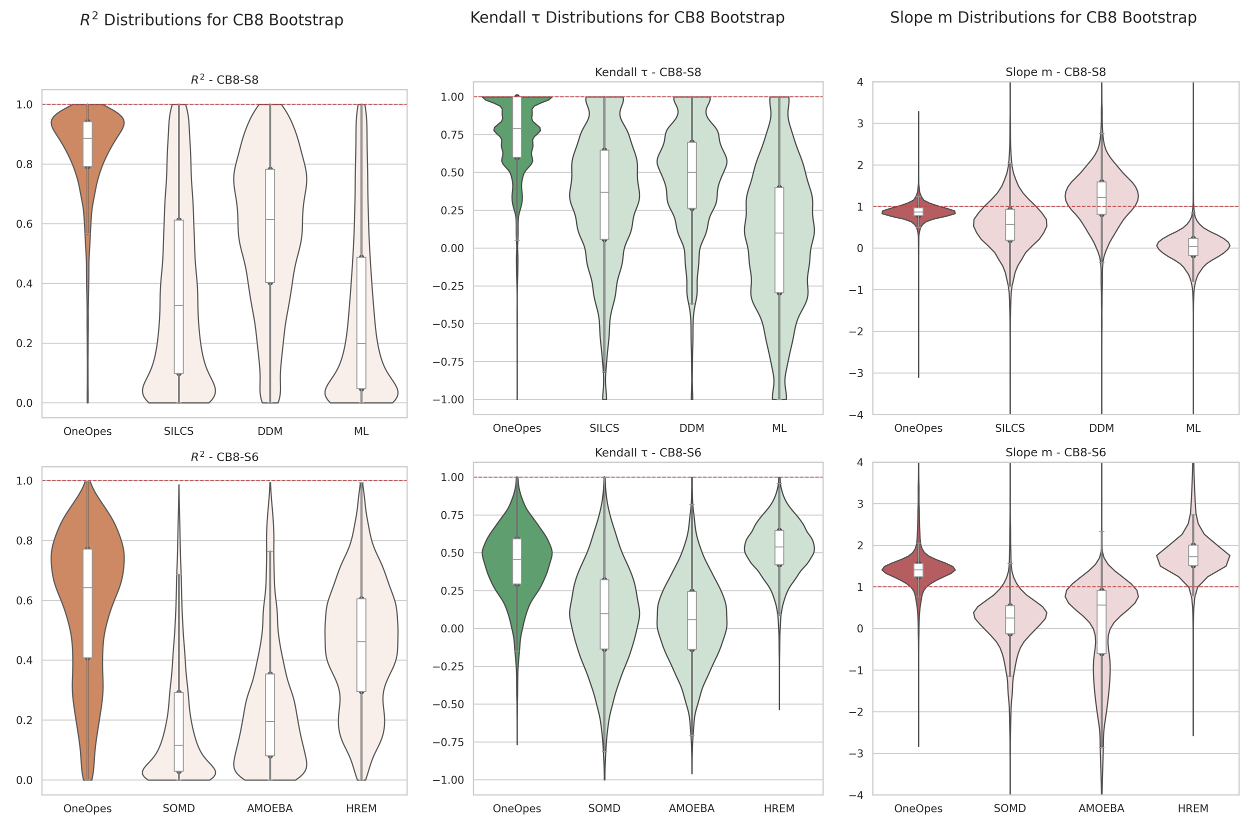


**Figure S1:** $R^2$, $\tau$ and $m$ metrics for host-guest systems with host CB8 belonging to the SAMPL8 (top row) and SAMPL6 (bottom row) challenges. Confidence intervals are calculated with a bootstrap procedure. We compare the metrics resulting from OneOpes with three of the best performing submissions from each SAMPL challenge. The red line in the plots indicate the ideal value of each metric. The results from SAMPL8 are taken from Ref. 4 and those from SAMPL6 from Ref. 5. SILCS corresponds to submission id 5, DDM[6] corresponds to submission id 28, ML[7] corresponds to submission id 10, SOMD corresponds to submission id 15, AMOEBA corresponds to submission id 33 and HREM corresponds to submission id 4.

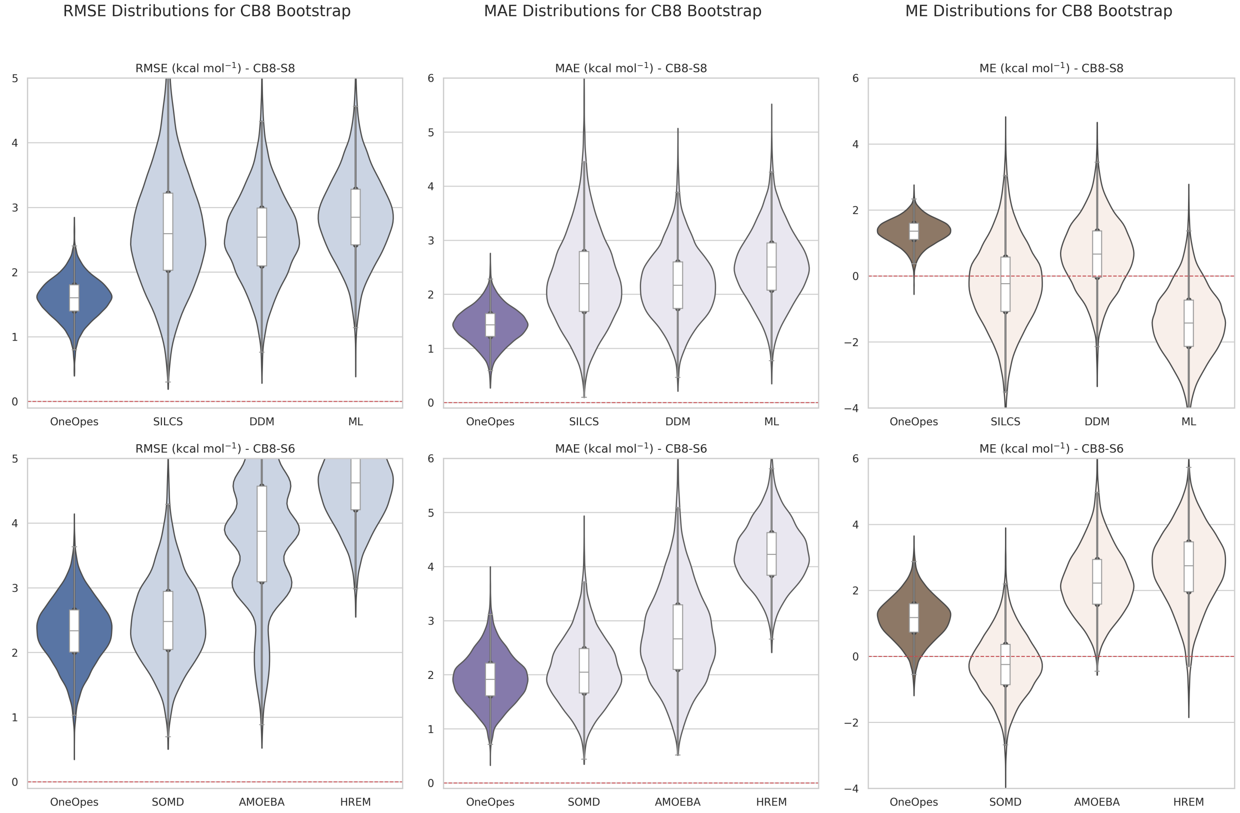


**Figure S2:** RMSE, MAE and ME metrics for host-guest systems with host CB8 belonging to the SAMPL8 (top row) and SAMPL6 (bottom row) challenges. Confidence intervals are calculated with a bootstrap procedure. We compare the metrics resulting from OneOpes with three of the best performing submissions from each SAMPL challenge. The red line in the plots indicate the ideal value of each metric. The results from SAMPL8 are taken from Ref. 4 and those from SAMPL6 from Ref. 5. SILCS corresponds to submission id 5, DDM[6] corresponds to submission id 28, ML[7] corresponds to submission id 10, SOMD corresponds to submission id 15, AMOEBA corresponds to submission id 33 and HREM corresponds to submission id 4.

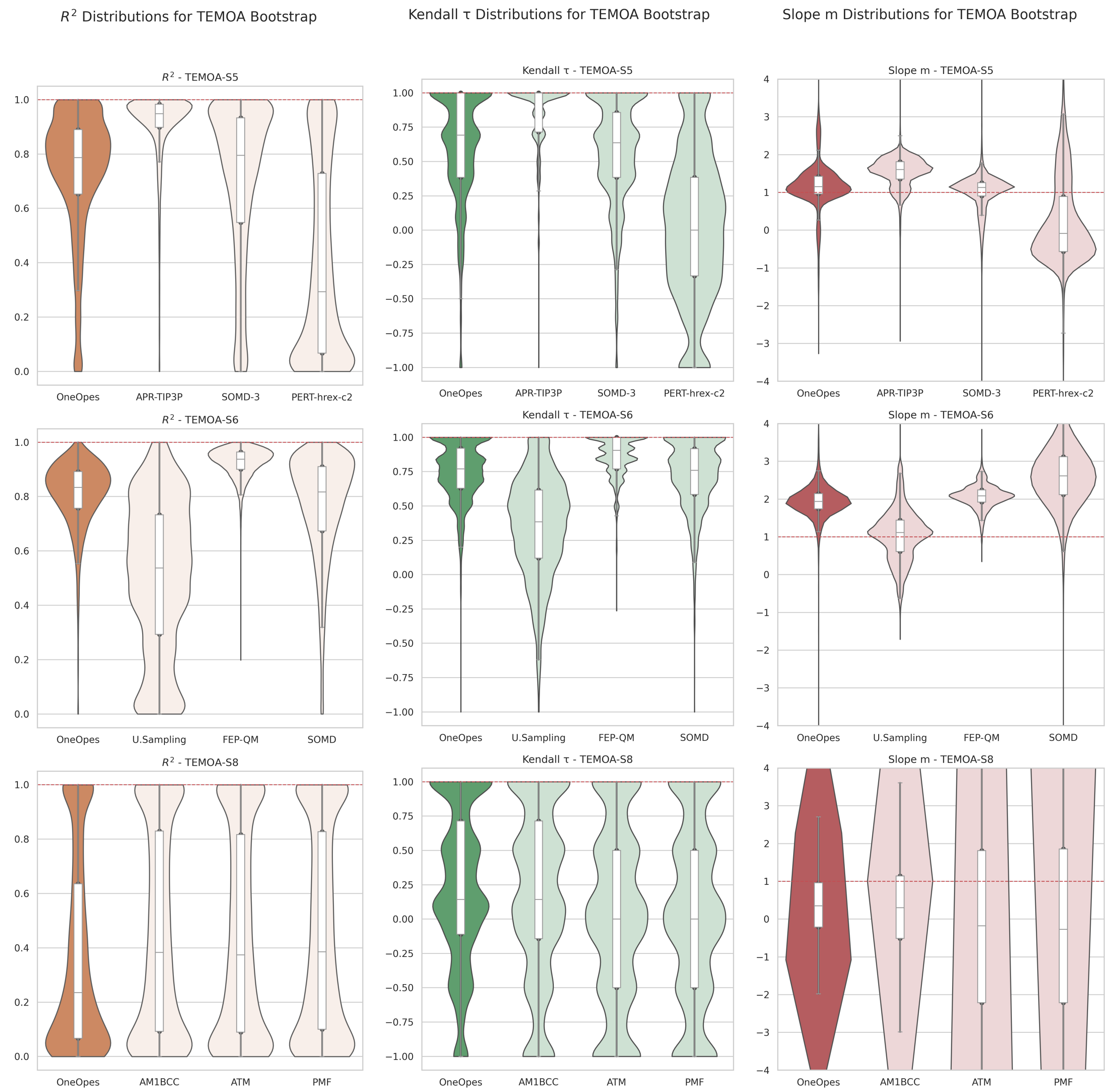


**Figure S3:** $R^2$, $\tau$ and $m$ metrics for host-guest systems with host TEMOA belonging to the SAMPL5 (top row), SAMPL6 (middle row) and SAMPL8 (bottom row) challenges. Confidence intervals are calculated with a bootstrap procedure. We compare the metrics resulting from OneOpes with three of the best performing submissions from each SAMPL challenge. The red line in the plots indicate the ideal value of each metric. The results from SAMPL8 are taken from Ref. 4, the results from SAMPL6 from Ref. 5 and the results from SAMPL5 from Ref. 8. AM1BCC corresponds to submission id 12, ATM[9] corresponds to submission id 1, PMF[9] corresponds to submission id 13, U.Sampling corresponds to submission id 39, FEP-QM corresponds to submission id 35, SOMD corresponds to submission id 21, APR-TIP3P[10] corresponds to submission id 2, SOMD-3[11] corresponds to submission id 24 and PERT-hrex-c2[12] corresponds to submission id 21.

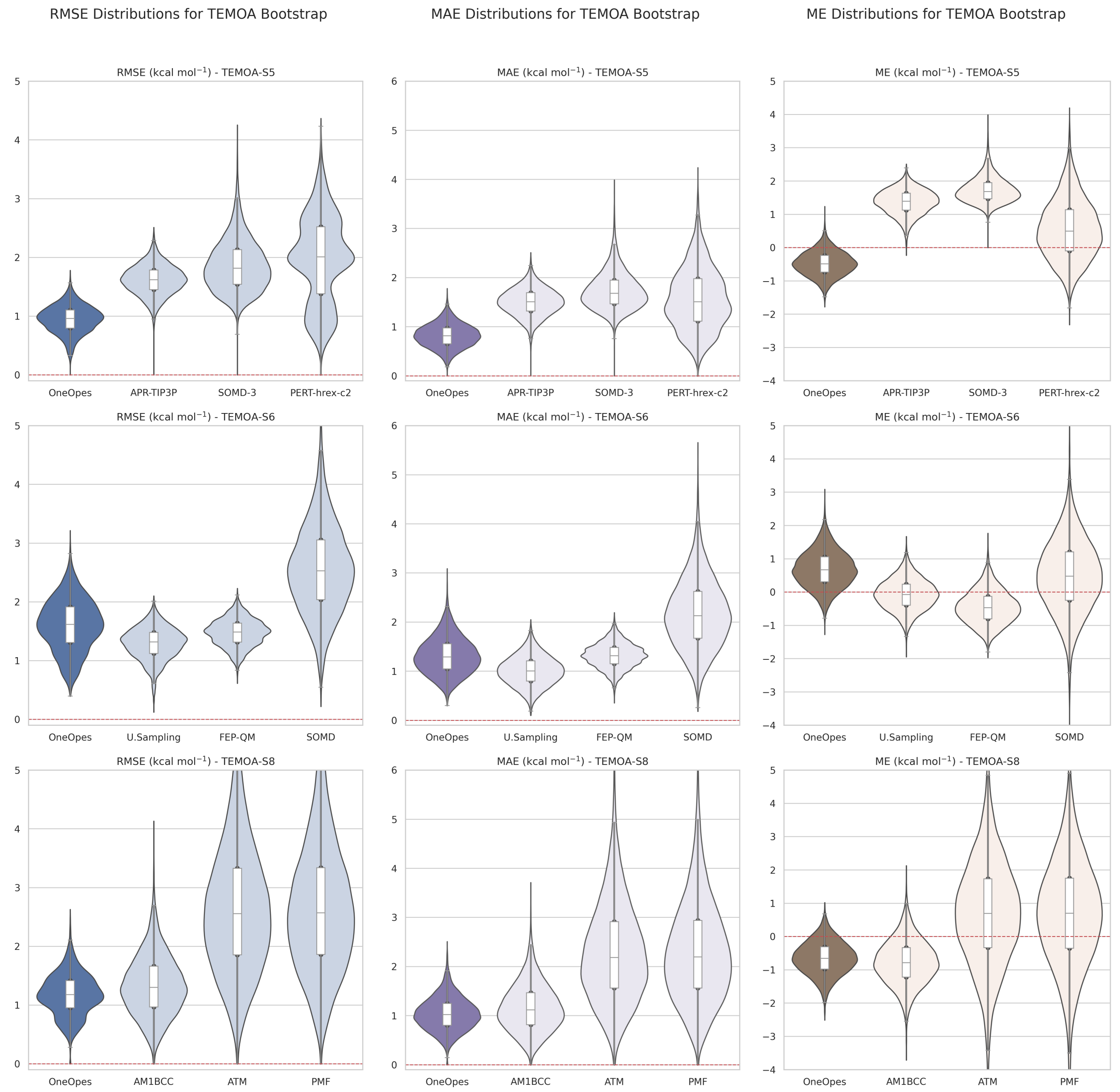


**Figure S4:** RMSE, MAE and ME metrics for host-guest systems with host TEMOA belonging to the SAMPL5 (top row), SAMPL6 (middle row) and SAMPL8 (bottom row) challenges. Confidence intervals are calculated with a bootstrap procedure. We compare the metrics resulting from OneOpes with three of the best performing submissions from each SAMPL challenge. The red line in the plots indicate the ideal value of each metric. The results from SAMPL8 are taken from Ref. 4, the results from SAMPL6 from Ref. 5 and the results from SAMPL5 from Ref. 8. AM1BCC corresponds to submission id 12, ATM[9] corresponds to submission id 1, PMF[9] corresponds to submission id 13, U.Sampling corresponds to submission id 39, FEP-QM corresponds to submission id 35, SOMD corresponds to submission id 21, APR-TIP3P[10] corresponds to submission id 2, SOMD-3[11] corresponds to submission id 24 and PERT-hrex-c2[12] corresponds to submission id 21.

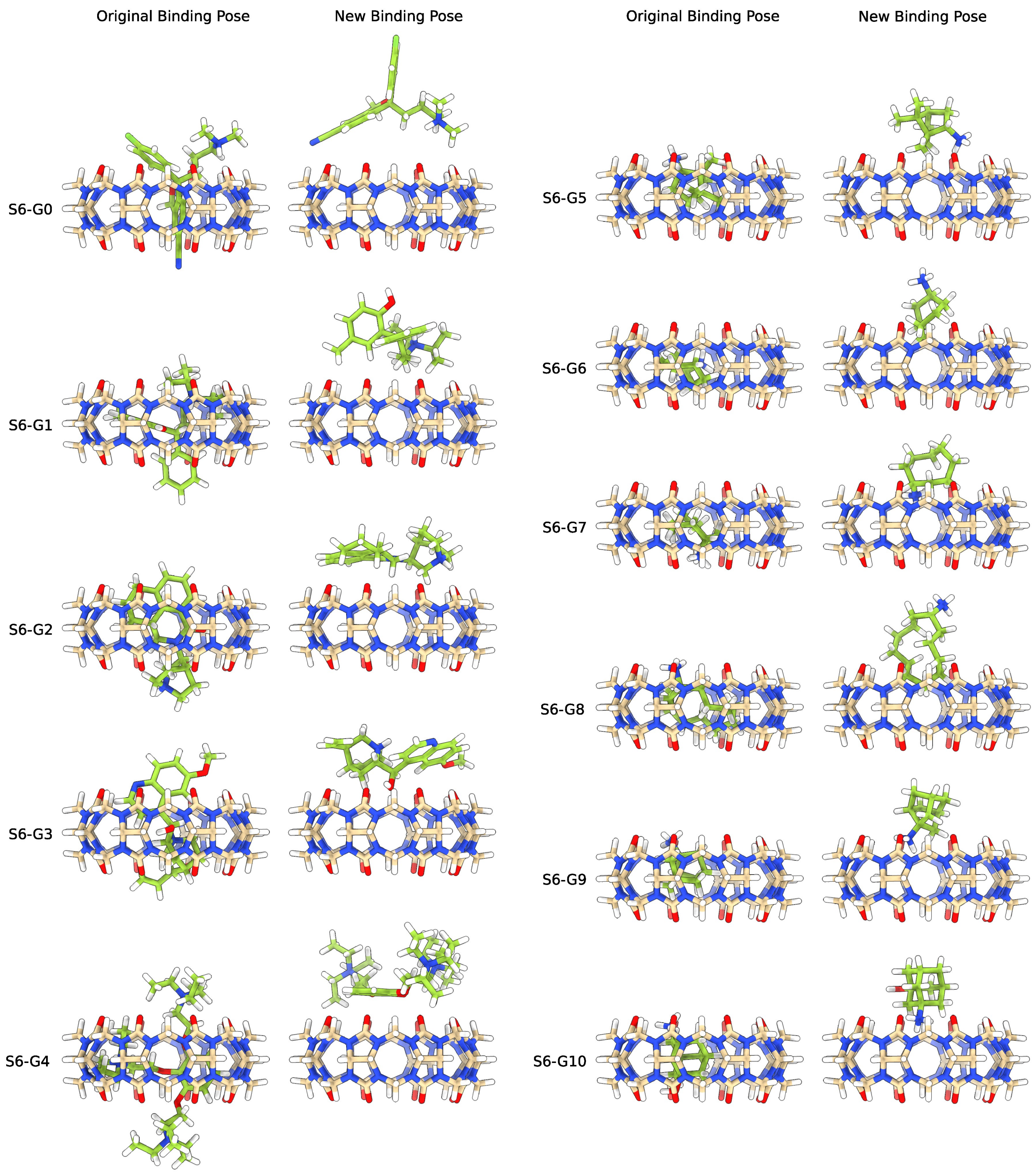


**Figure S5:** Initial binding poses that we simulated for host CB8 and guests belonging to the SAMPL6 challenge. On the left hand side, we show the original binding pose provided by the SAMPL challenge organisers. On the right hand side, we show the alternative initial state that we tested that often differs significantly from the expected binding pose.

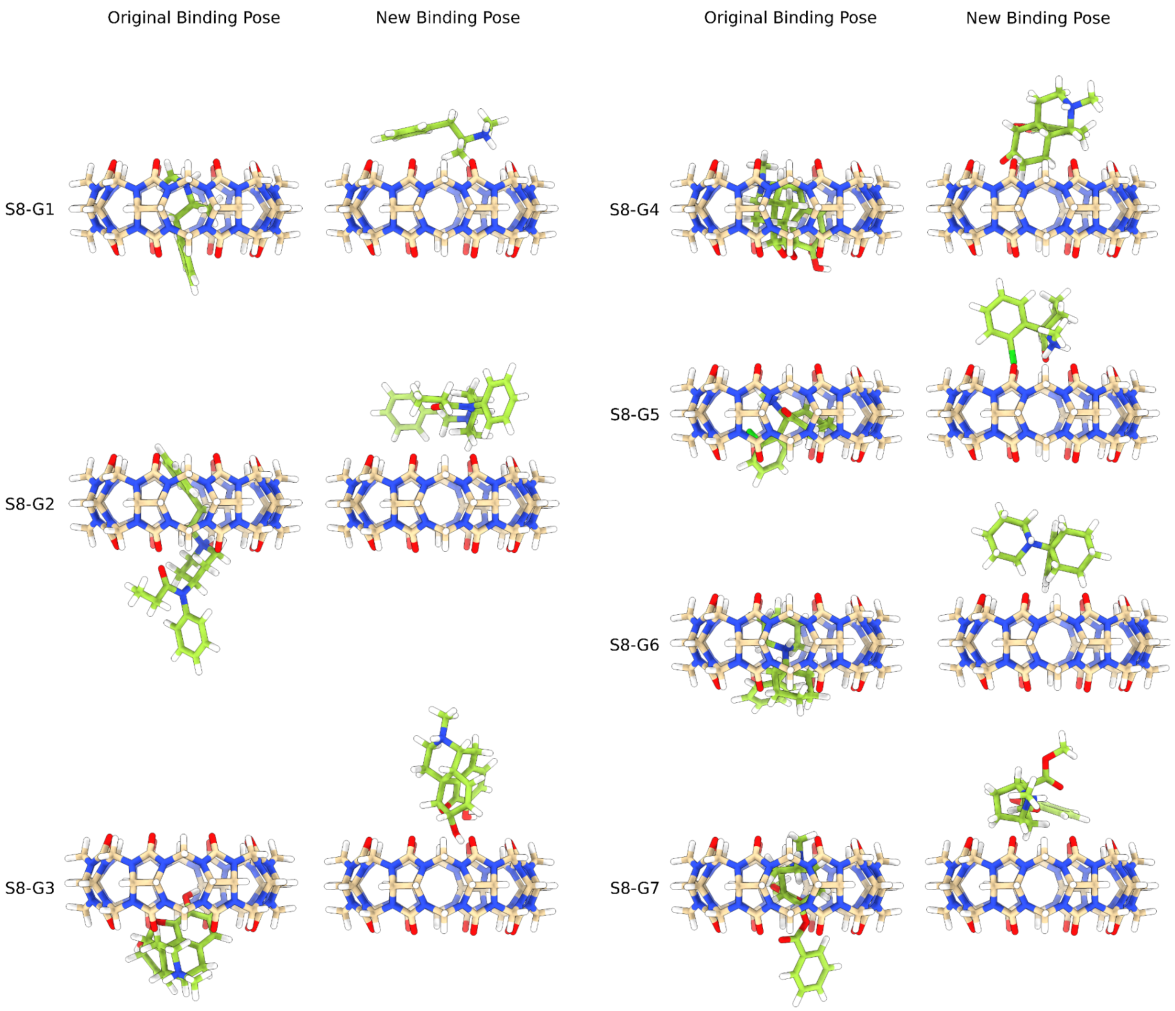


**Figure S6:** Initial binding poses that we simulated for host CB8 and guests belonging to the SAMPL6 challenge. On the left hand side, we show the original binding pose provided by the SAMPL challenge organisers. On the right hand side, we show the alternative initial state that we tested that often differs significantly from the expected binding pose.

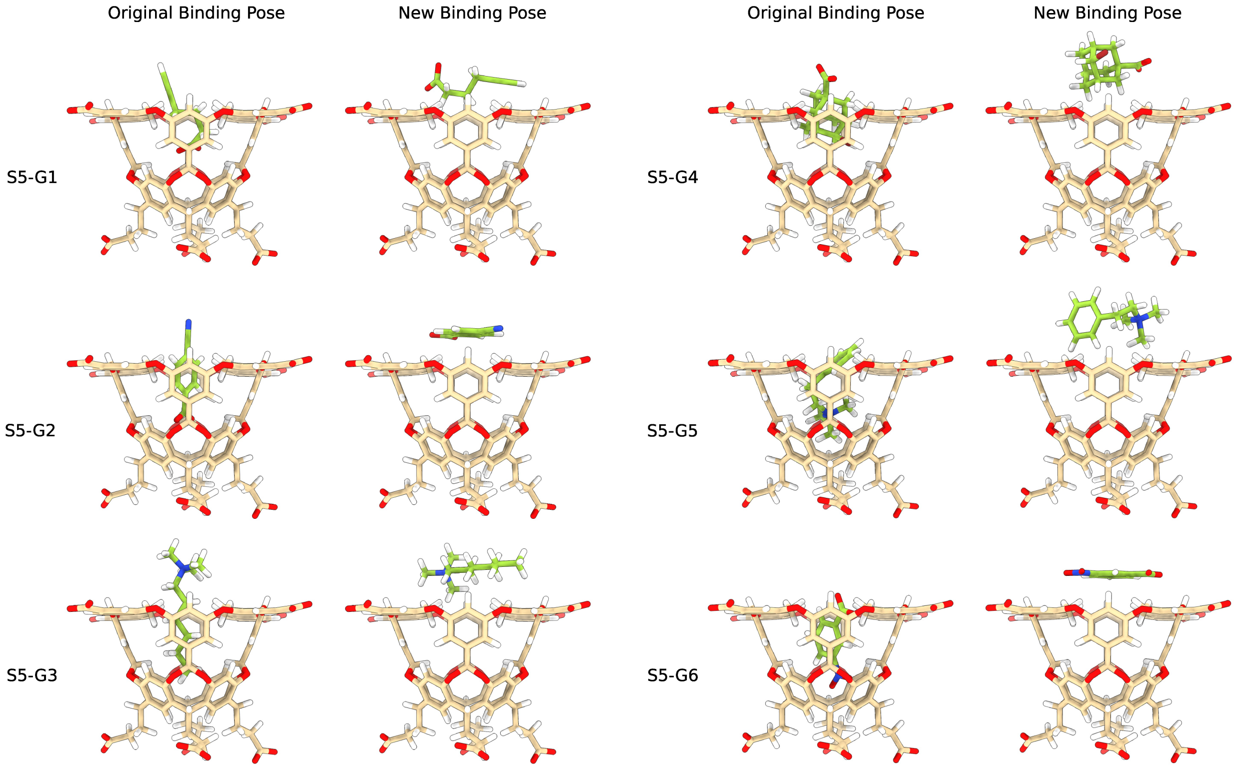


**Figure S7:** Initial binding poses that we simulated for host TEMOA and guests belonging to the SAMPL5 challenge. On the left hand side, we show the original binding pose provided by the SAMPL challenge organisers. On the right hand side, we show the alternative initial state that we tested that often differs significantly from the expected binding pose.

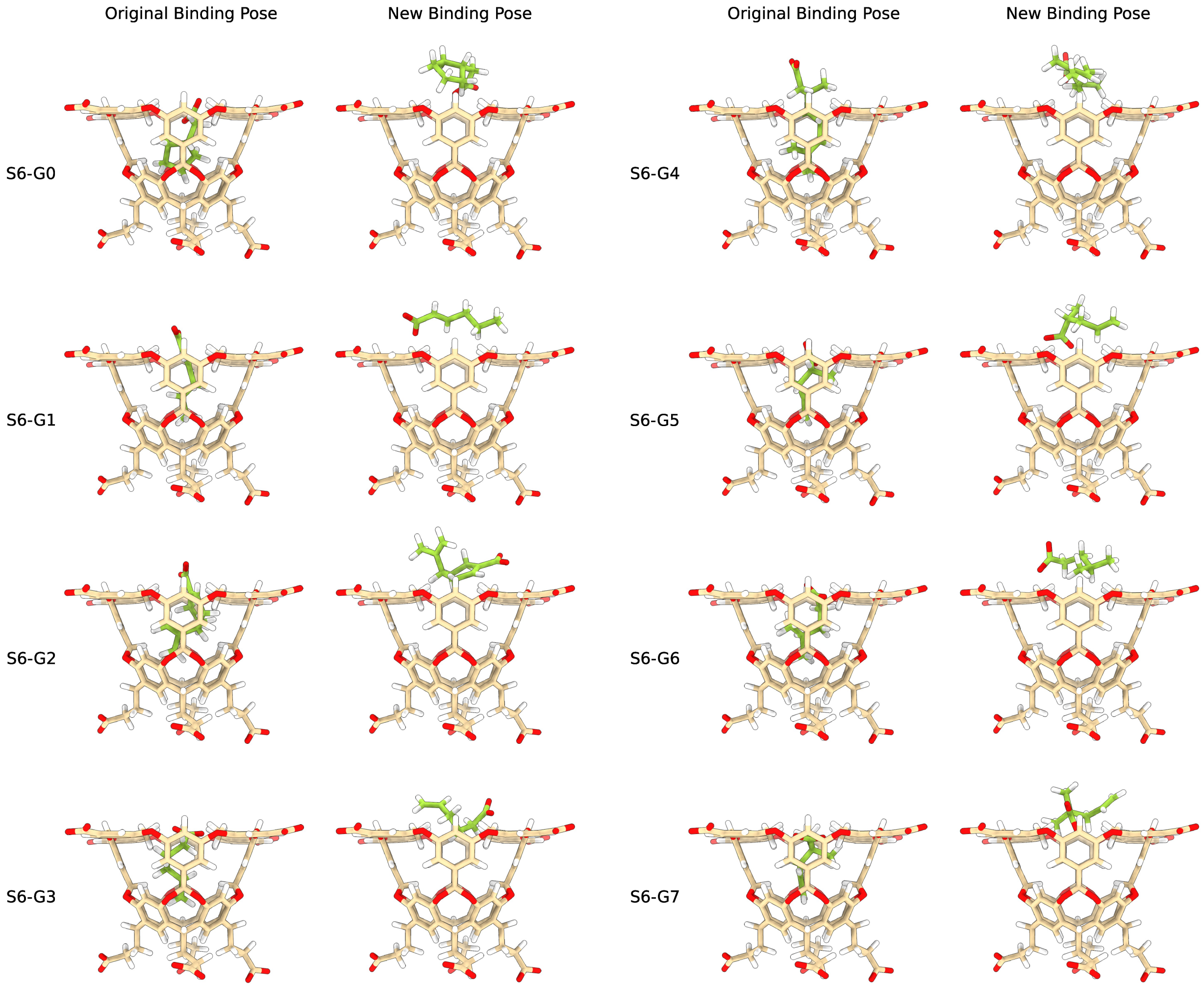


**Figure S8:** Initial binding poses that we simulated for host TEMOA and guests belonging to the SAMPL6 challenge. On the left hand side, we show the original binding pose provided by the SAMPL challenge organisers. On the right hand side, we show the alternative initial state that we tested that often differs significantly from the expected binding pose.

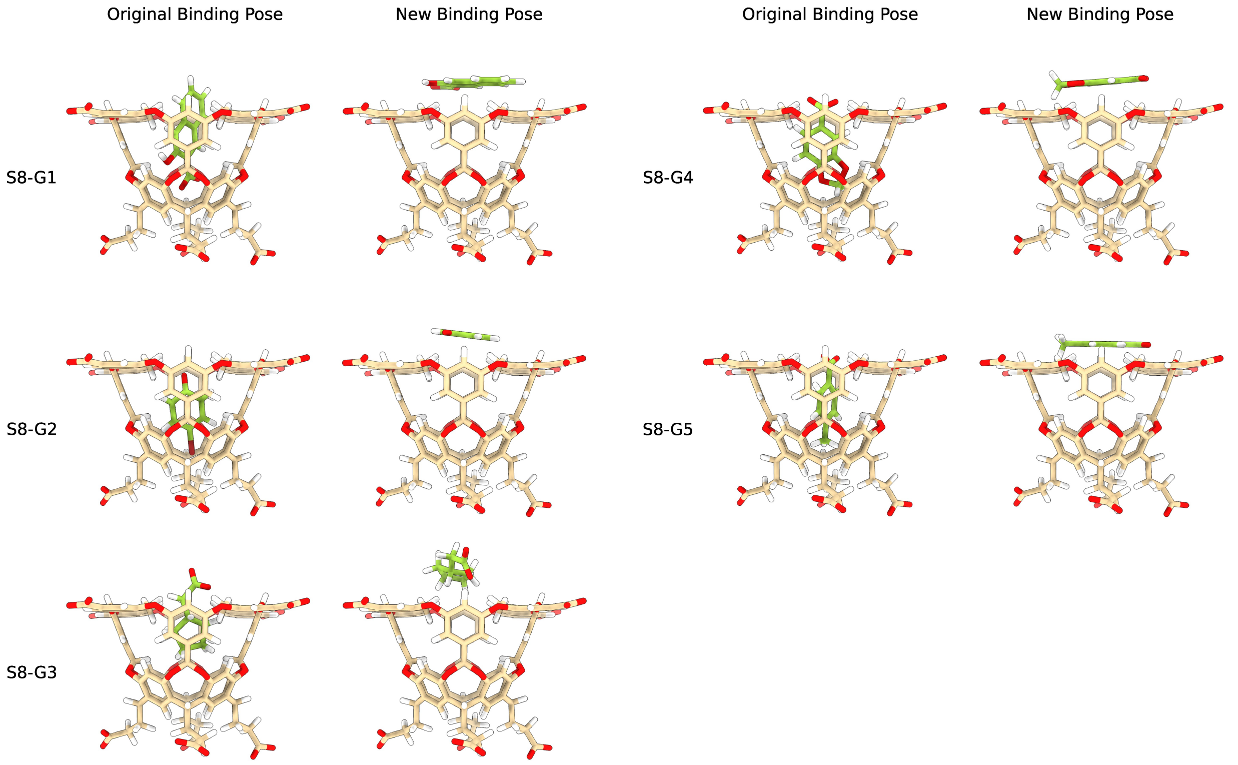


**Figure S9:** Initial binding poses that we simulated for host TEMOA and guests belonging to the SAMPL8 challenge. On the left hand side, we show the original binding pose provided by the SAMPL challenge organisers. On the right hand side, we show the alternative initial state that we tested that often differs significantly from the expected binding pose.

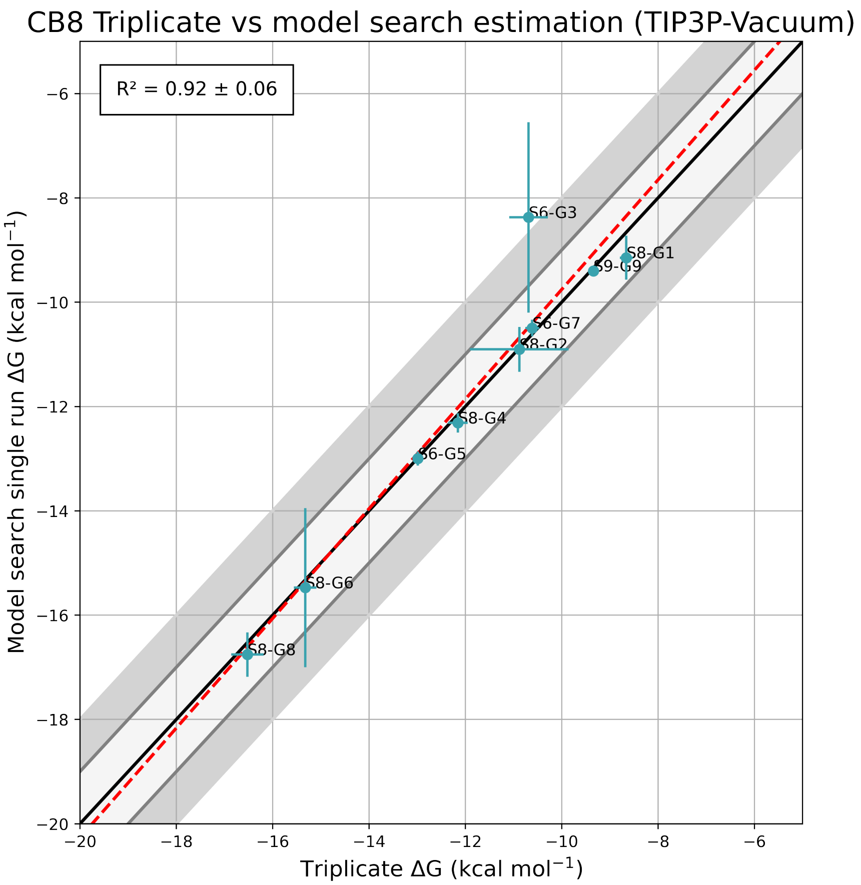


**Figure S10:** Correlation plot comparing the binding free energies obtained from the Force Field Search phase and the triplicate simulations from the Result Refinement phase of host CB8. The force field used includes TIP3P water and vacuum electrostatics. The solid black line represents the line of perfect correlation, and the red dashed line is the best-fit linear regression. The corresponding $R^2 = 0.92 \pm 0.06$ indicates a strong correlation between the two sets of data points.

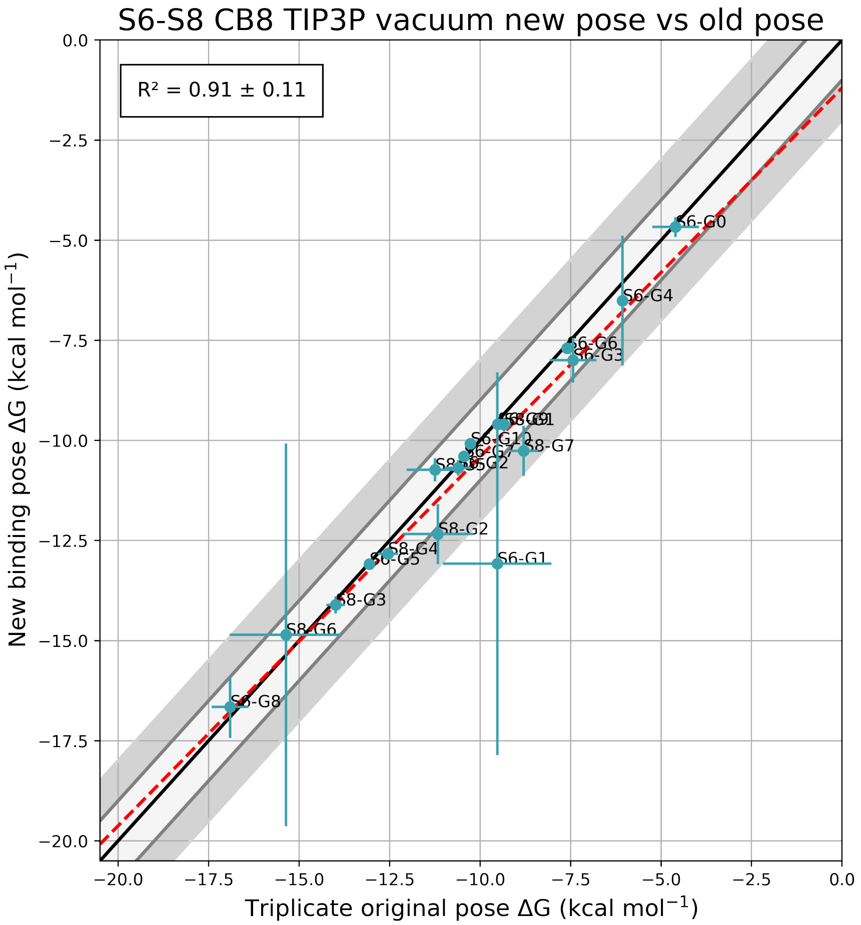


**Figure S11:** Correlation plot comparing the binding free energies obtained from the triplicate simulations from the Result Refinement phase and another triplicate set of simulations starting from an alternative initial state in host CB8. The force field used includes TIP3P water and vacuum electrostatics. The solid black line represents the line of perfect correlation, and the red dashed line is the best-fit linear regression. The corresponding $R^2 = 0.91 \pm 0.11$ indicates a strong correlation between the the original binding pose and the alternative initial state.

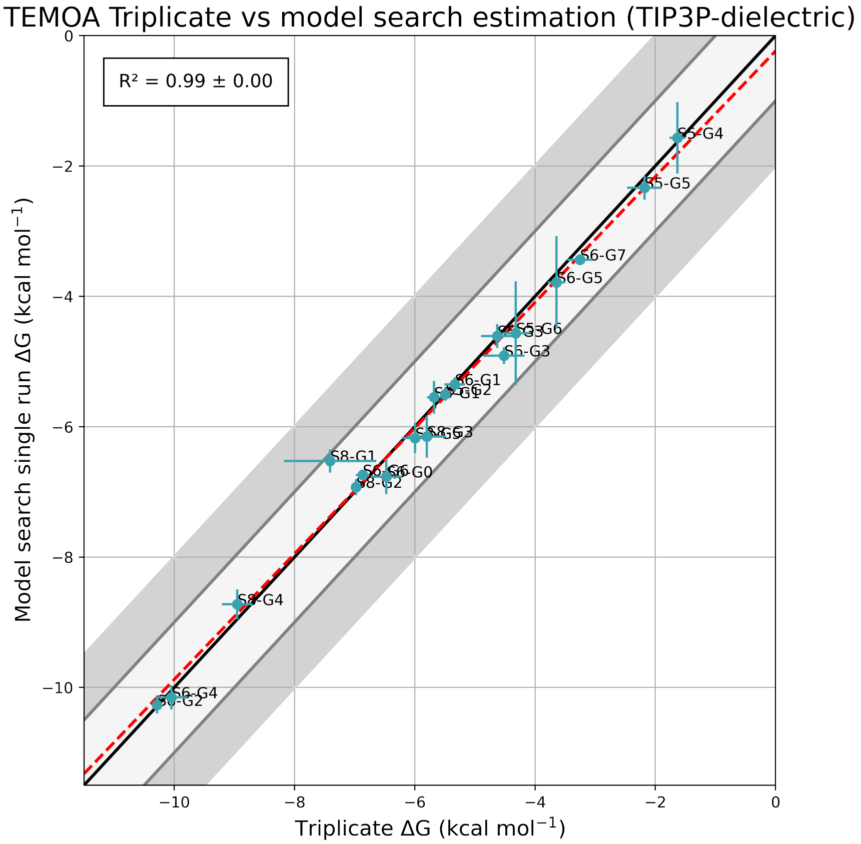


**Figure S12:** Correlation plot comparing the binding free energies obtained from the Force Field Search phase and the triplicate simulations from the Result Refinement phase of host TEMOA. The force field used includes TIP3P water and dielectric electrostatics. The solid black line represents the line of perfect correlation, and the red dashed line is the best-fit linear regression. The corresponding $R^2 = 0.99 \pm 0.06$ indicates a strong correlation between the two sets of data points.

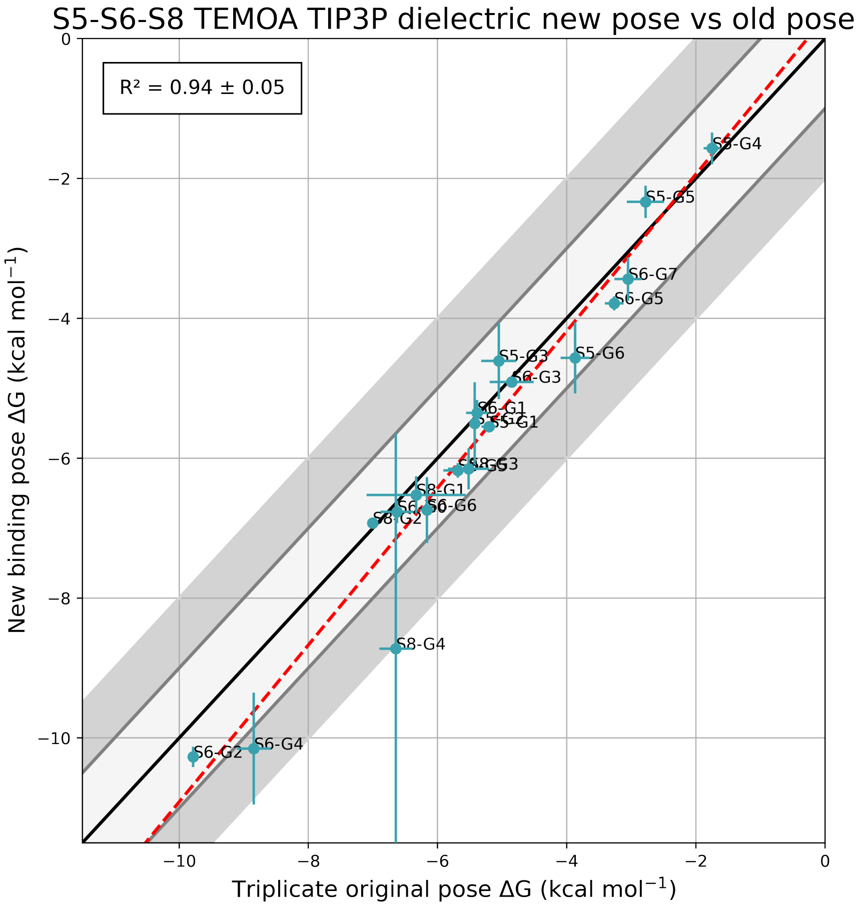


**Figure S13:** Correlation plot comparing the binding free energies obtained from the triplicate simulations from the Result Refinement phase and another triplicate set of simulations starting from an alternative initial state in host TEMOA. The force field used includes TIP3P water and dielectric electrostatics. The solid black line represents the line of perfect correlation, and the red dashed line is the best-fit linear regression. The corresponding $R^2 = 0.94 \pm 0.05$ indicates a strong correlation between the the original binding pose and the alternative initial state.

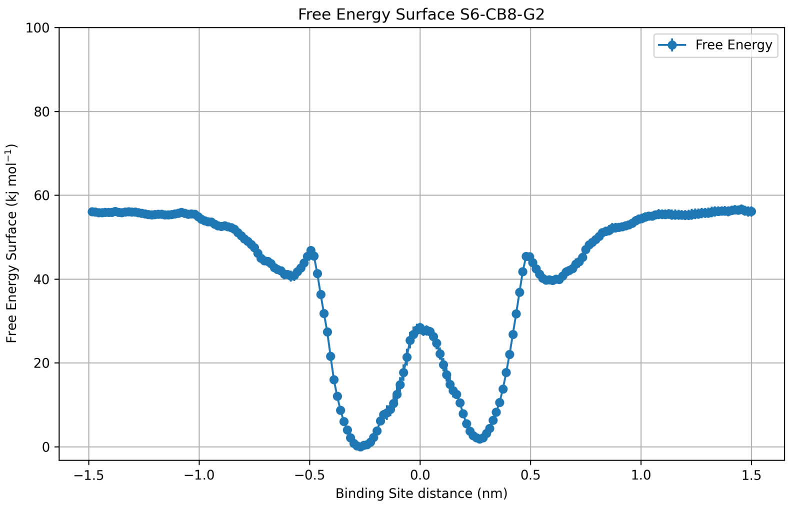


**Figure S14:** Free energy profile as a function of $z$ of the system with host CB8 and guest S6-G2. Two minima are present. The apparent symmetry of the free energy profiles is not imposed and corresponds perfectly to the symmetry of the host molecule. This shows how well it has converged.

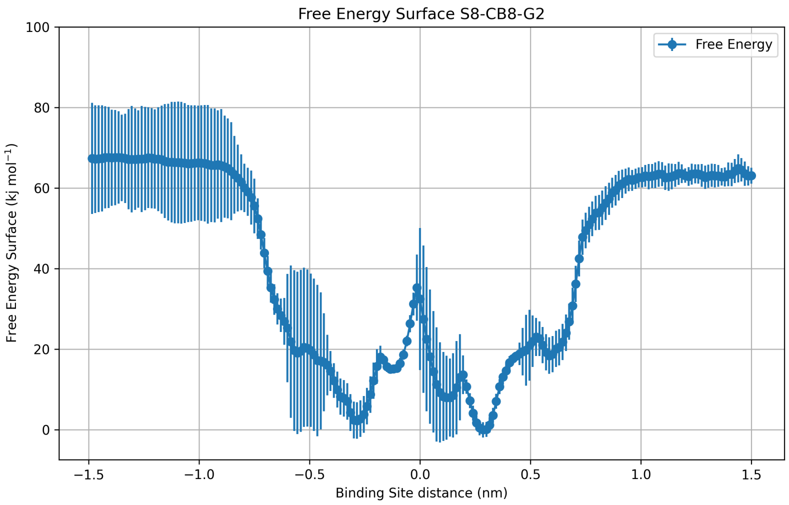


**Figure S15:** Free energy profile as a function of $z$ of the system with host CB8 and guest S8-G2. As the guest is rather large and flexible, the free energy profile is more noisy. It presents a rich binding landscape and preserves a good degree of symmetry between the upper and lower funnel.

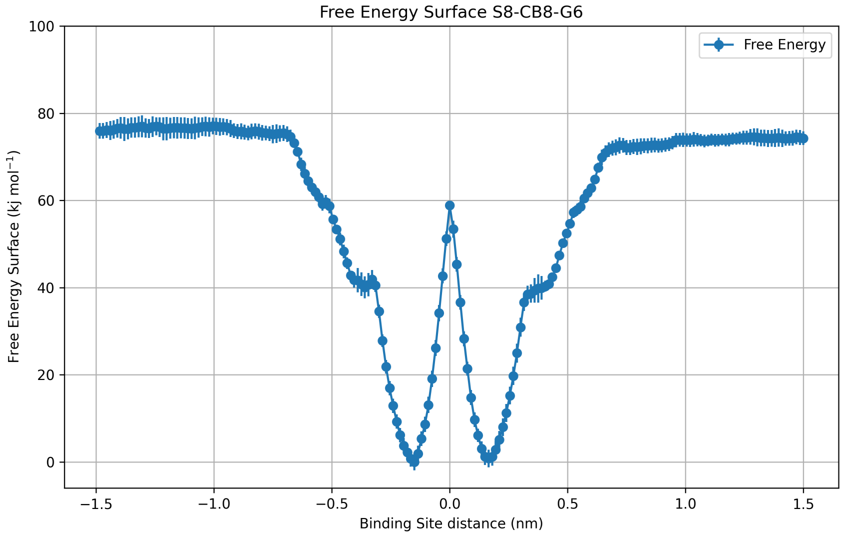


**Figure S16:** Free energy profile as a function of $z$ of the system with host CB8 and guest S8-G6. Two closely packed minima are visible.

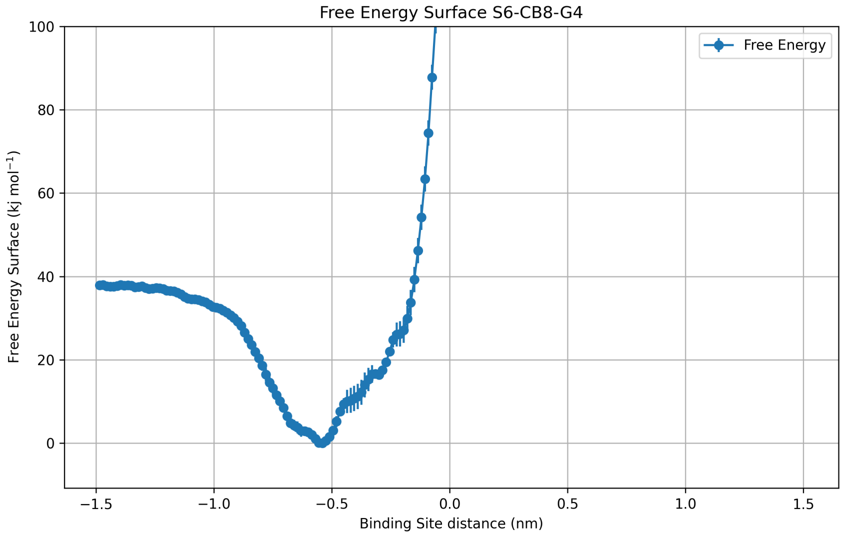


**Figure S17:** Free energy profile as a function of $z$ of the system with host CB8 and guest S6-G4. This large ligand is not able to squeeze through the host, hence it samples well only one of the two binding sites.

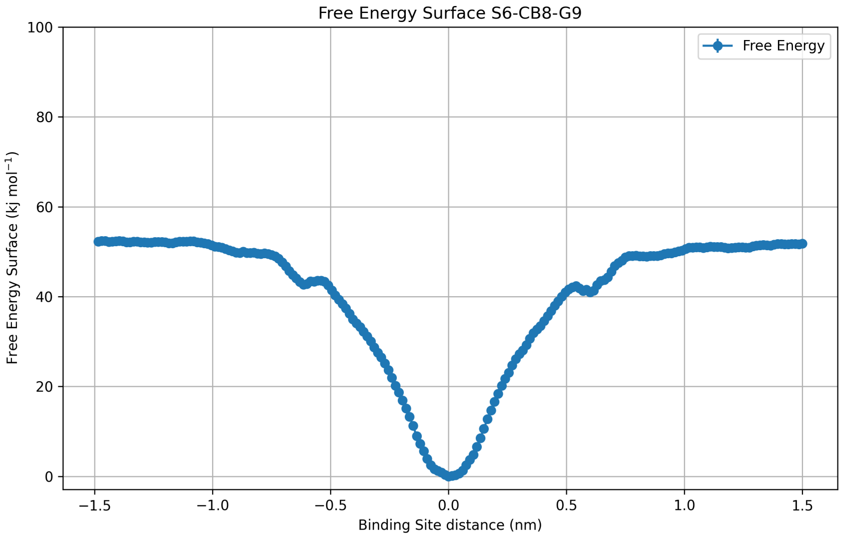


**Figure S18:** Free energy profile as a function of $z$ of the system with host CB8 and guest S 6-G9. A single minimum is present and the profile displays a good symmetry.

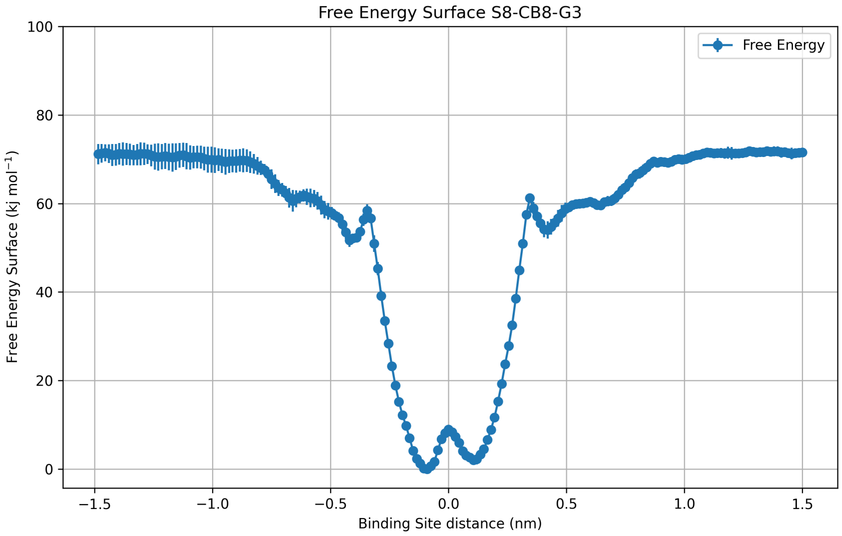


**Figure S19:** Free energy profile as a function of $z$ of the system with host CB8 and guest S8-G3. Two closely packed minima are visible and the profile displays a good symmetry.

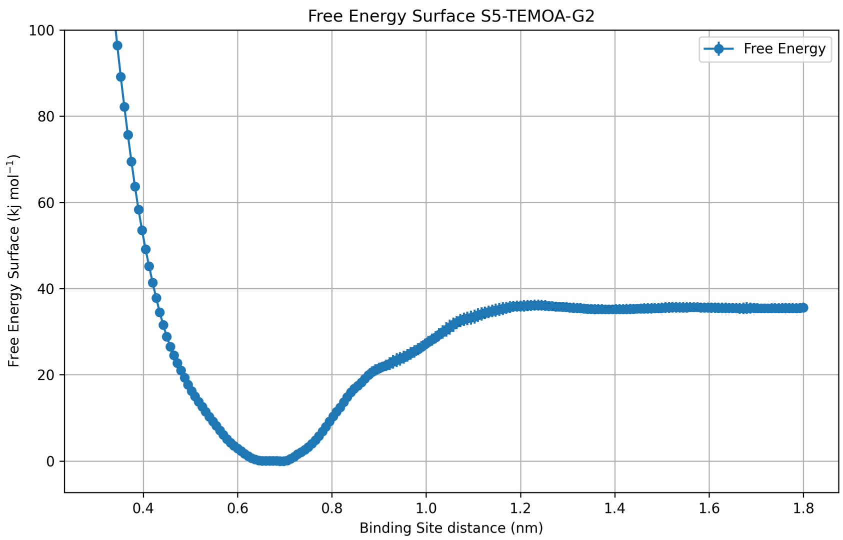


**Figure S20:** Free energy profile as a function of $z$ of the system with host TEMOA and guest S5-G2.

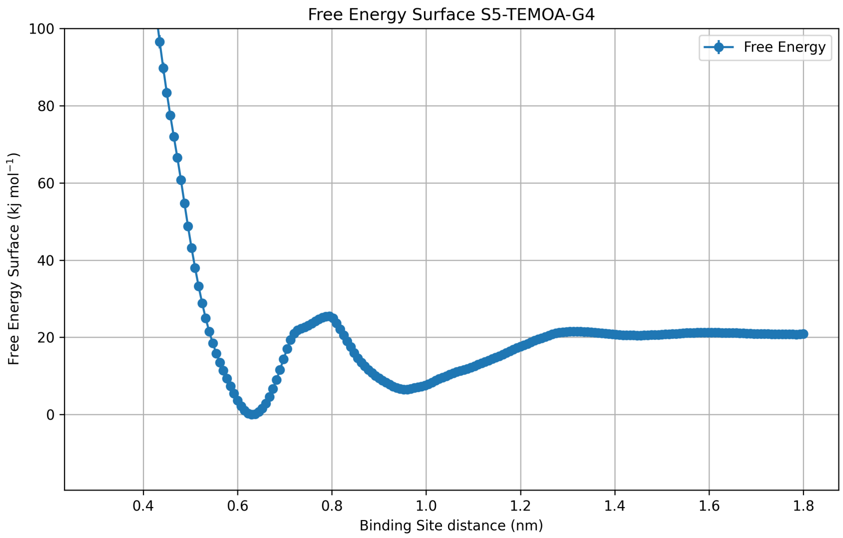


**Figure S21:** Free energy profile as a function of $z$ of the system with host TEMOA and guest S5-G4.

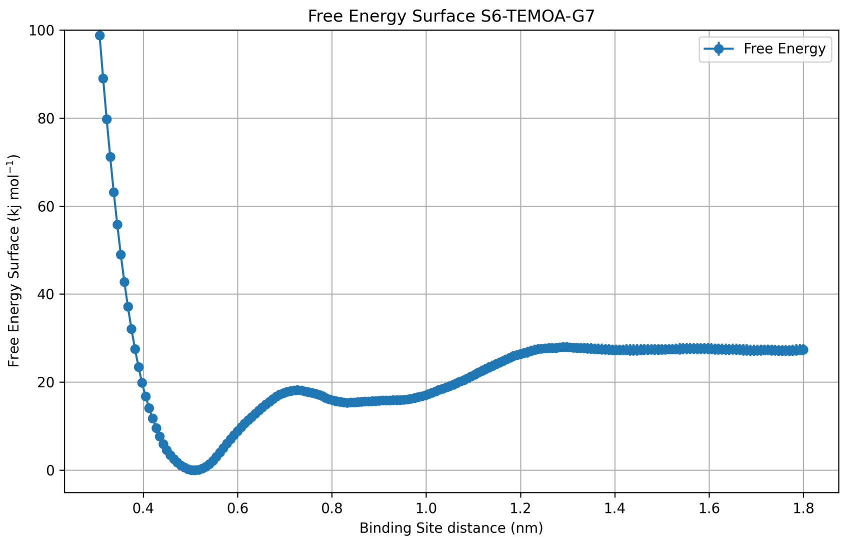


**Figure S22:** Free energy profile as a function of $z$ of the system with host TEMOA and guest S6-G7.

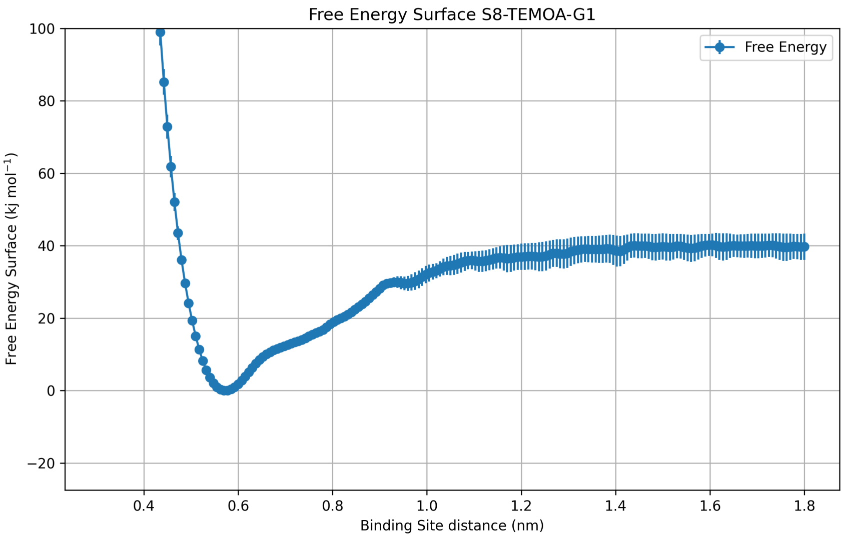


**Figure S23:** Free energy profile as a function of $z$ of the system with host TEMOA and guest S8-G1.

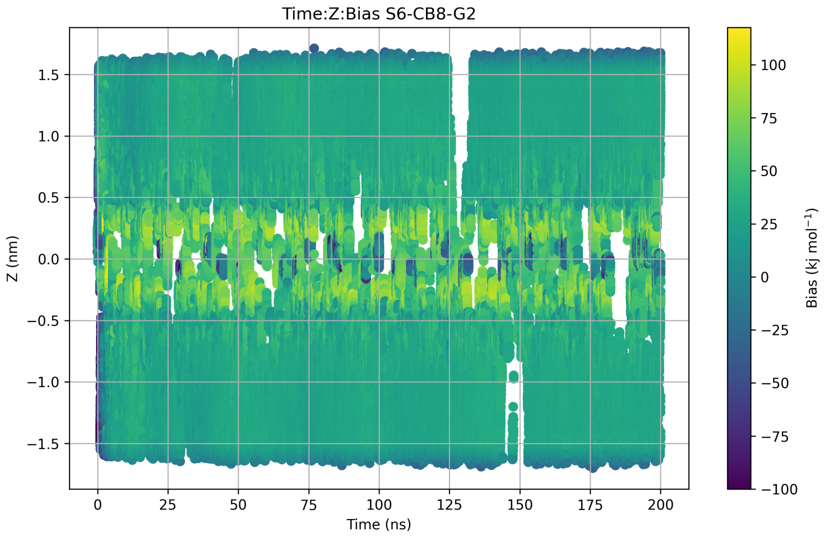


**Figure S24:** Dynamics of $z$ in replica 0 of a OneOPES simulation of host CB8 and guest S6-G2 from the Results Refinement phase. The plot is colored with the instantaneous value of the OPES Explore bias.

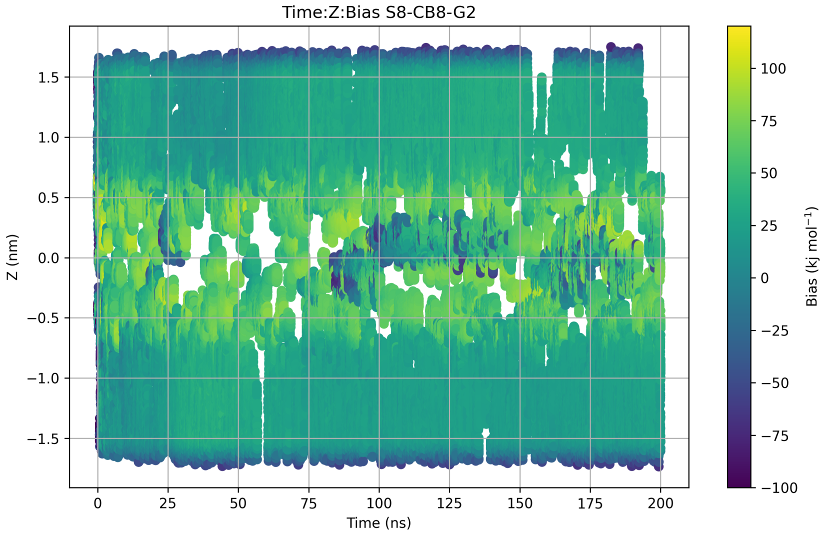


**Figure S25:** Dynamics of $z$ in replica 0 of a OneOPES simulation of host CB8 and guest S8-G2 from the Results Refinement phase. The plot is colored with the instantaneous value of the OPES Explore bias.

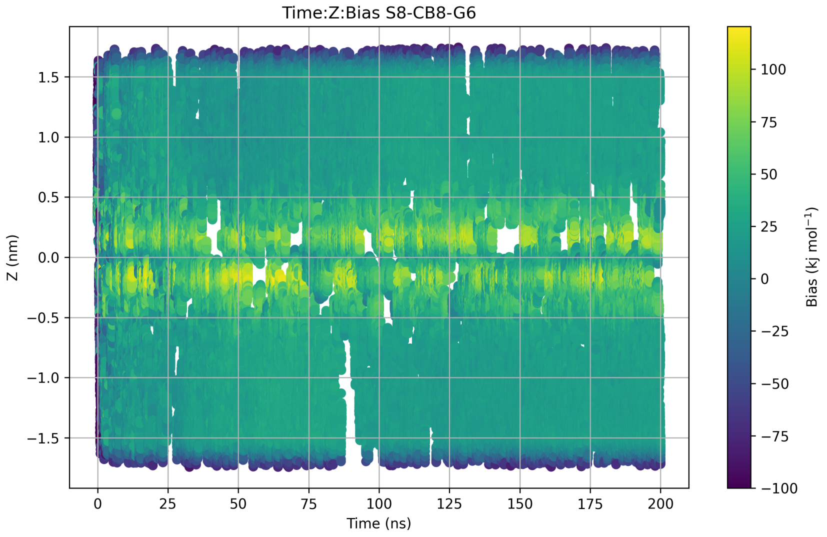


**Figure S26:** Dynamics of $z$ in replica 0 of a OneOPES simulation of host CB8 and guest S8-G6 from the Results Refinement phase. The plot is colored with the instantaneous value of the OPES Explore bias.

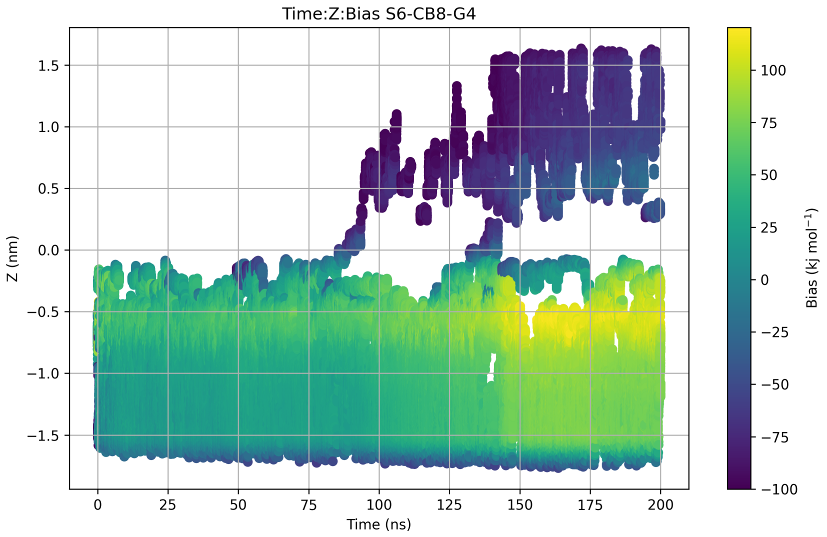


**Figure S27:** Dynamics of $z$ in replica 0 of a OneOPES simulation of host CB8 and guest S6-G4 from the Results Refinement phase. The plot is colored with the instantaneous value of the OPES Explore bias.

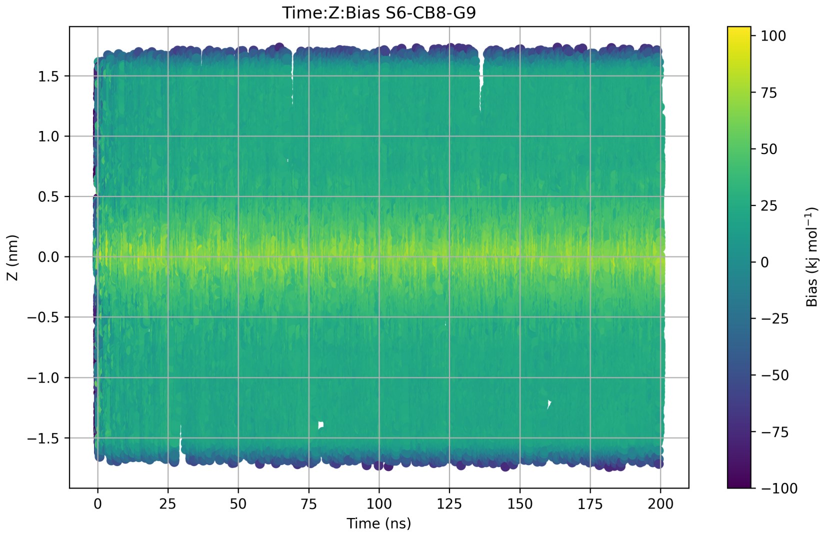


**Figure S28:** Dynamics of $z$ in replica 0 of a OneOPES simulation of host CB8 and guest S6-G9 from the Results Refinement phase. The plot is colored with the instantaneous value of the OPES Explore bias.

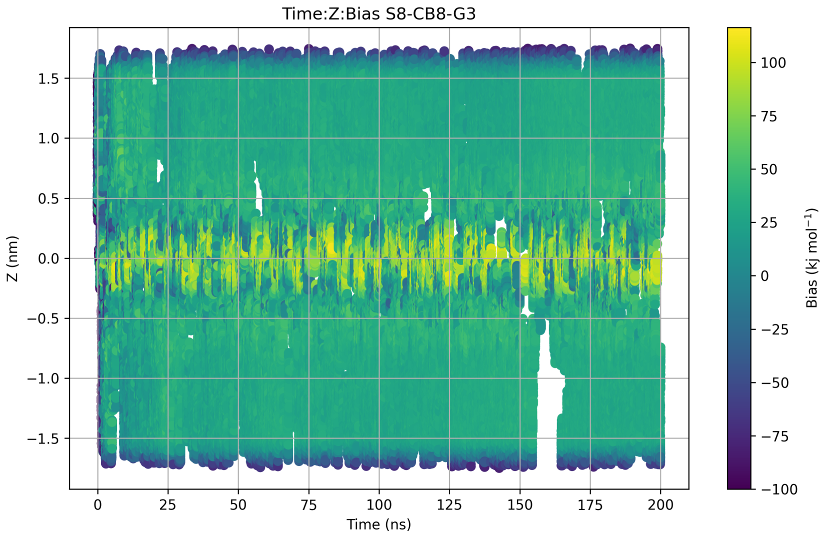


**Figure S29:** Dynamics of $z$ in replica 0 of a OneOPES simulation of host CB8 and guest S8-G3 from the Results Refinement phase. The plot is colored with the instantaneous value of the OPES Explore bias.

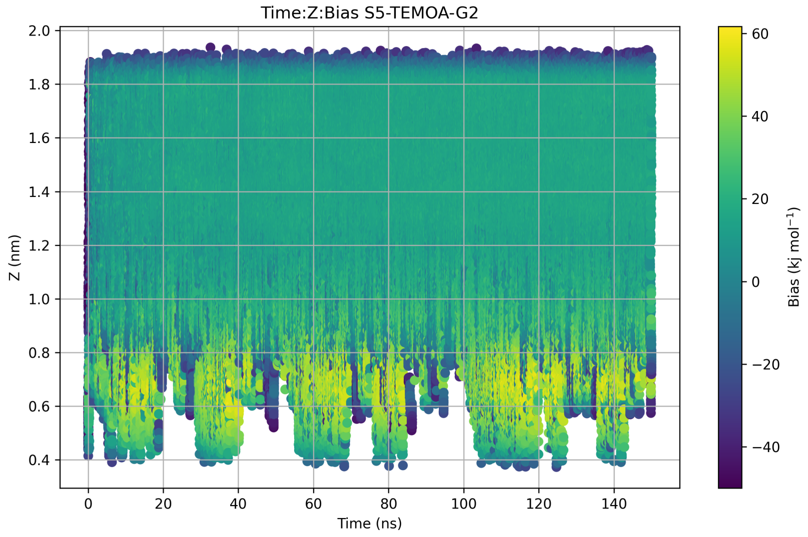


**Figure S30:** Dynamics of $z$ in replica 0 of a OneOPES simulation of host TEMOA and guest S5-G2 from the Results Refinement phase. The plot is colored with the instantaneous value of the OPES Explore bias.

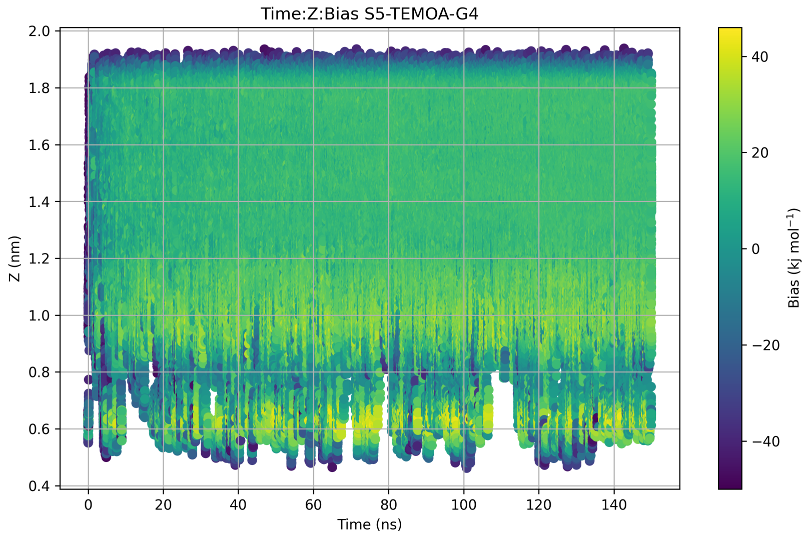


**Figure S31:** Dynamics of $z$ in replica 0 of a OneOPES simulation of host TEMOA and guest S5-G4 from the Results Refinement phase. The plot is colored with the instantaneous value of the OPES Explore bias.

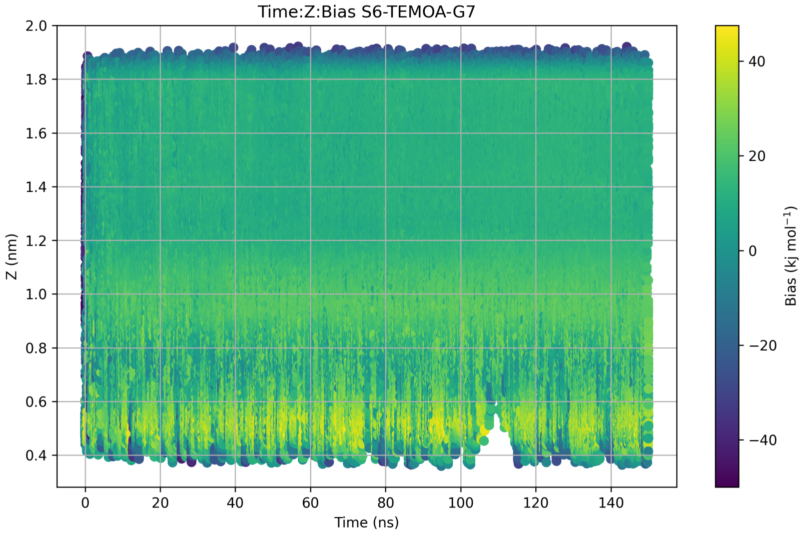


**Figure S32:** Dynamics of $z$ in replica 0 of a OneOPES simulation of host TEMOA and guest S6-G7 from the Results Refinement phase. The plot is colored with the instantaneous value of the OPES Explore bias.

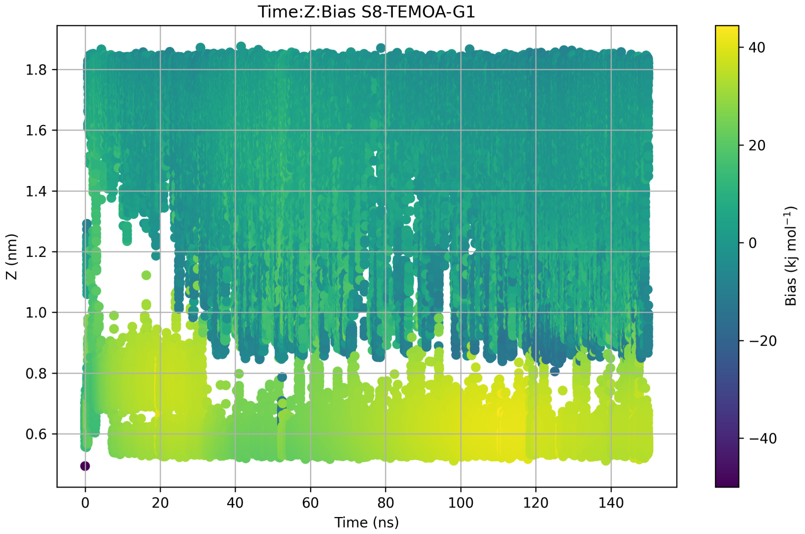


**Figure S33:** Dynamics of $z$ in replica 0 of a OneOPES simulation of host TEMOA and guest S8-G1 from the Results Refinement phase. The plot is colored with the instantaneous value of the OPES Explore bias.

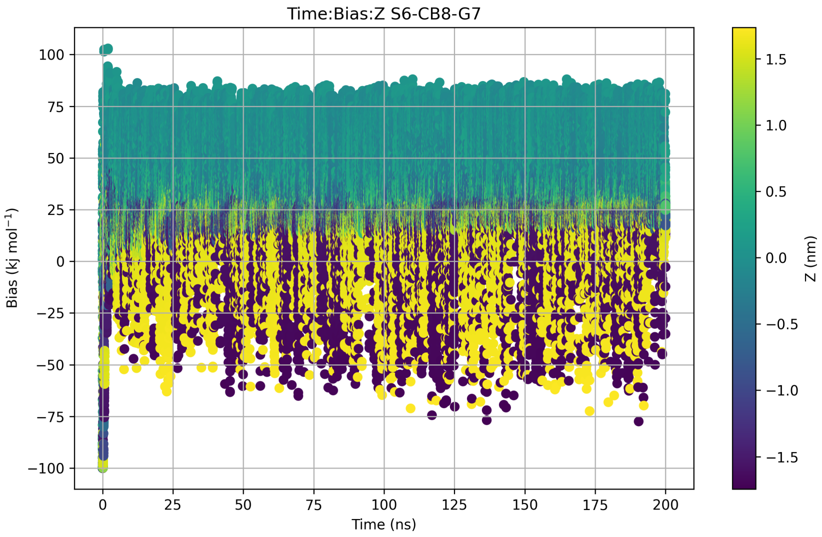


**Figure S34:** Dynamics of the OPES Explore bias in replica 0 of a OneOPES simulation of host CB8 and guest S6-G2 from the Results Refinement phase. The plot is colored with the instantaneous value of $z$.

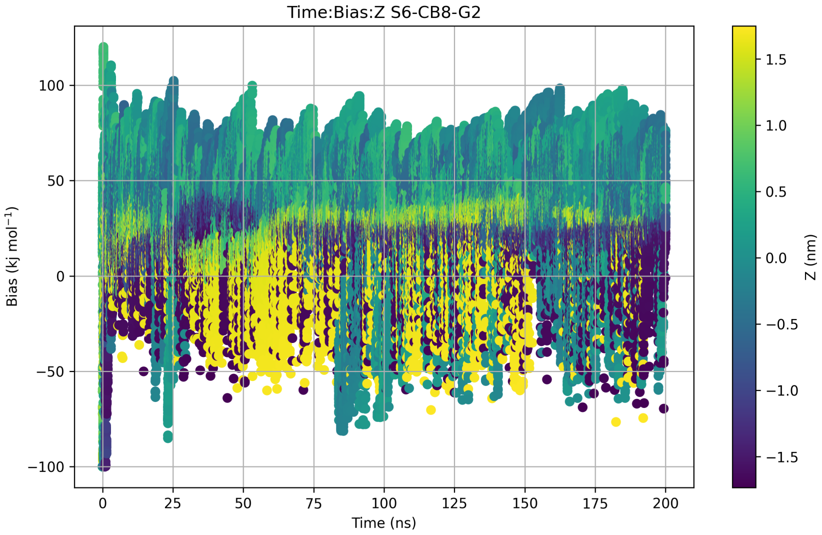


**Figure S35:** Dynamics of the OPES Explore bias in replica 0 of a OneOPES simulation of host CB8 and guest S8-G2 from the Results Refinement phase. The plot is colored with the instantaneous value of $z$.

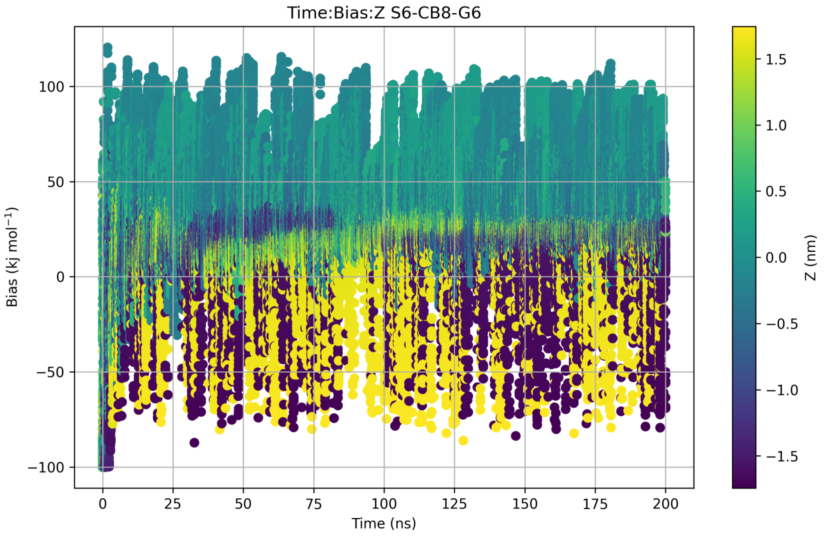


**Figure S36:** Dynamics of the OPES Explore bias in replica 0 of a OneOPES simulation of host CB8 and guest S8-G6 from the Results Refinement phase. The plot is colored with the instantaneous value of $z$.

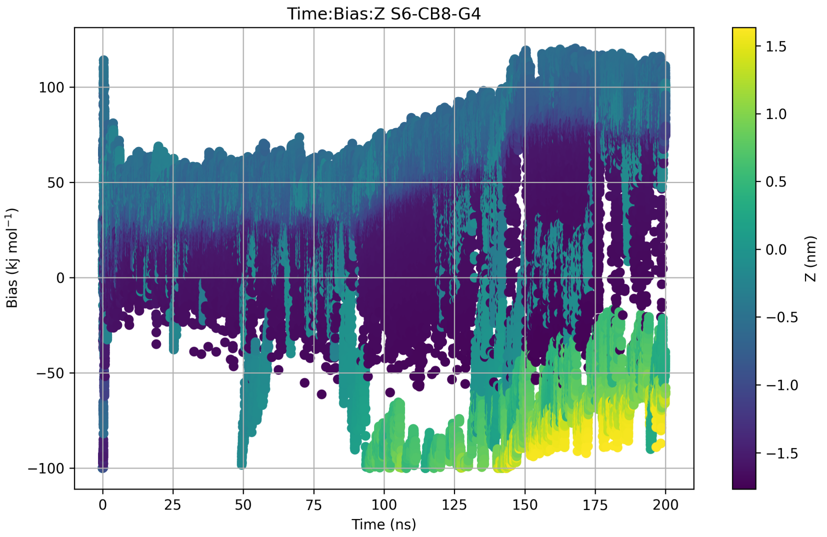


**Figure S37:** Dynamics of the OPES Explore bias in replica 0 of a OneOPES simulation of host CB8 and guest S6-G4 from the Results Refinement phase. The plot is colored with the instantaneous value of $z$.

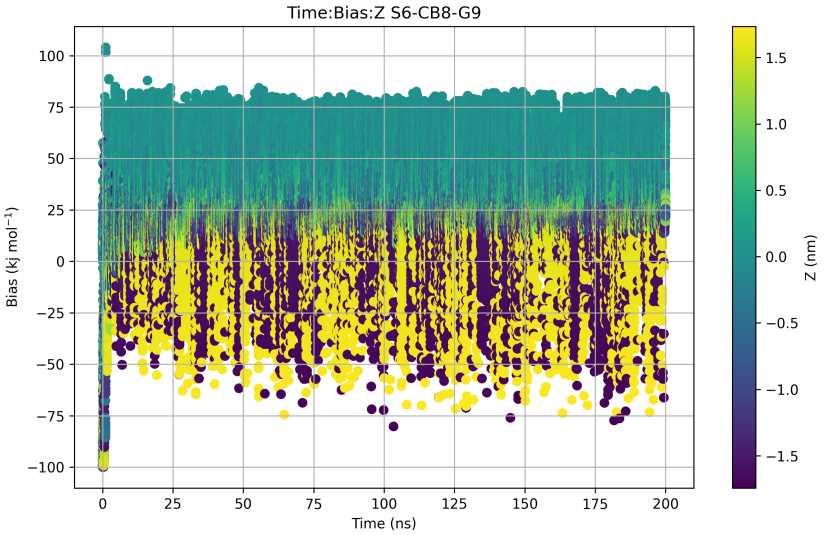


**Figure S38:** Dynamics of the OPES Explore bias in replica 0 of a OneOPES simulation of host CB8 and guest S6-G9 from the Results Refinement phase. The plot is colored with the instantaneous value of $z$.

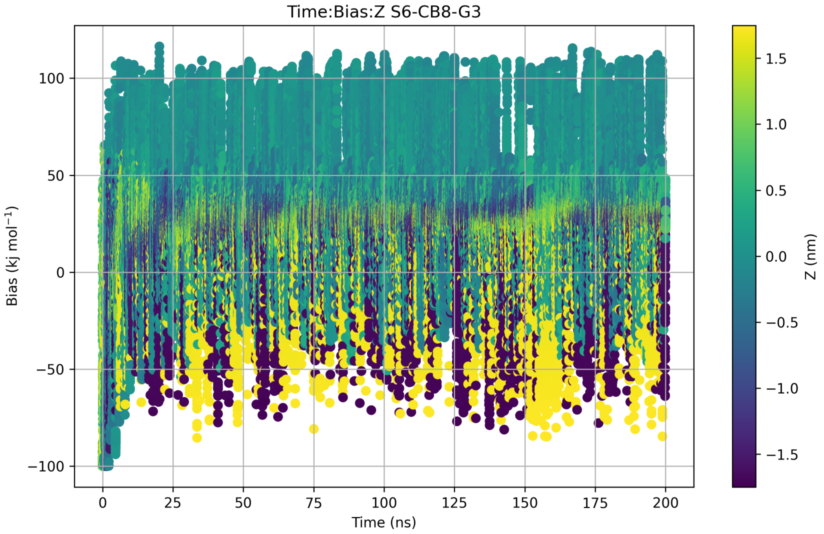


**Figure S39:** Dynamics of the OPES Explore bias in replica 0 of a OneOPES simulation of host CB8 and guest S8-G3 from the Results Refinement phase. The plot is colored with the instantaneous value of $z$.

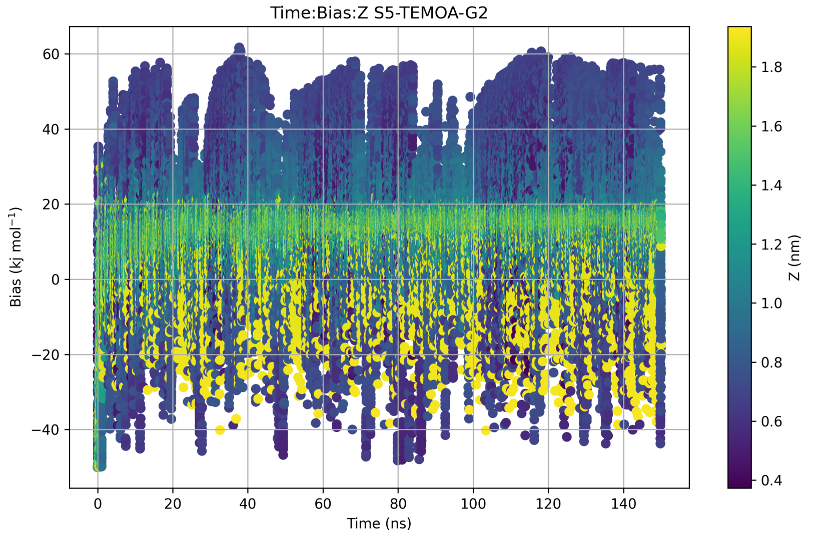


**Figure S40:** Dynamics of the OPES Explore bias in replica 0 of a OneOPES simulation of host TEMOA and guest S5-G2 from the Results Refinement phase. The plot is colored with the instantaneous value of $z$.

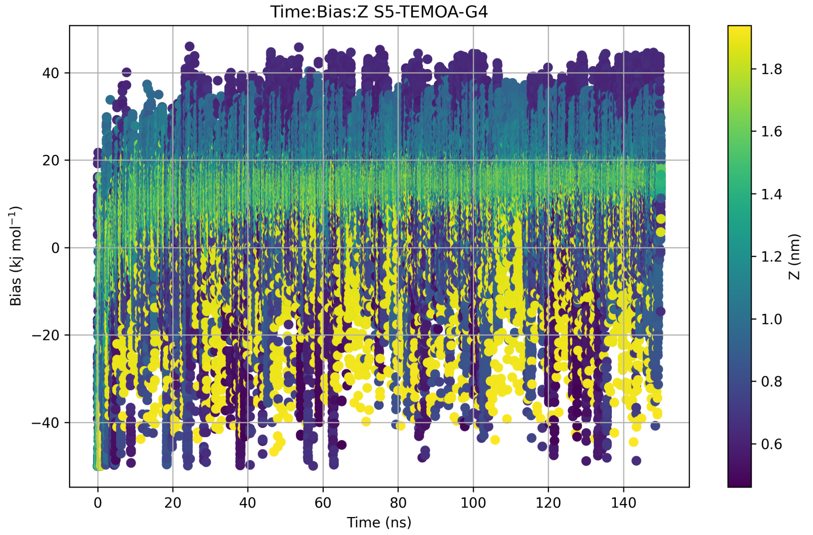


**Figure S41:** Dynamics of the OPES Explore bias in replica 0 of a OneOPES simulation of host TEMOA and guest S5-G4 from the Results Refinement phase. The plot is colored with the instantaneous value of $z$.

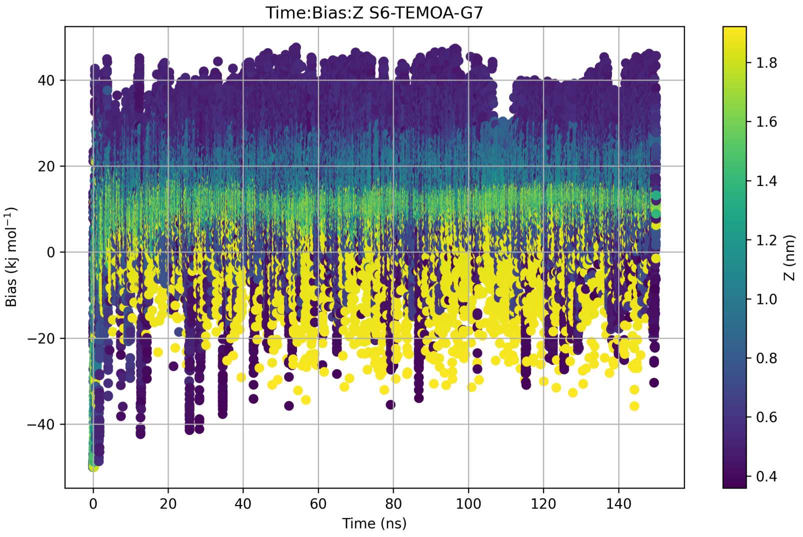


**Figure S42:** Dynamics of the OPES Explore bias in replica 0 of a OneOPES simulation of host TEMOA and guest S6-G7 from the Results Refinement phase. The plot is colored with the instantaneous value of $z$.

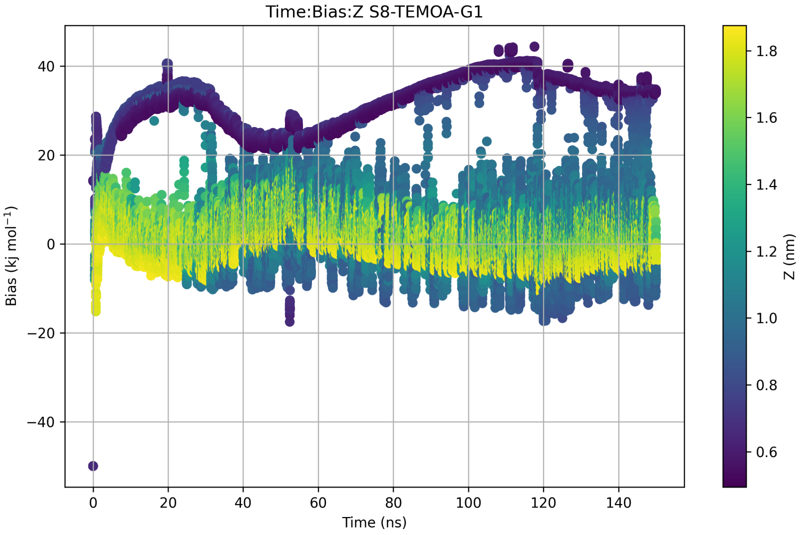


**Figure S43:** Dynamics of the OPES Explore bias in replica 0 of a OneOPES simulation of host TEMOA and guest S8-G1 from the Results Refinement phase. The plot is colored with the instantaneous value of $z$.
